# Synchrony between daily rhythms of malaria parasites and hosts is driven by an essential amino acid

**DOI:** 10.1101/2020.08.24.264689

**Authors:** Kimberley F. Prior, Benita Middleton, Alíz T.Y. Owolabi, Mary L. Westwood, Jacob Holland, Aidan J. O’Donnell, Mike Blackman, Debra J. Skene, Sarah E. Reece

## Abstract

Rapid asexual replication of blood stage malaria parasites is responsible for the severity of disease symptoms and fuels the production of transmission forms. That malaria parasite species coordinate their cycles of asexual replication with daily rhythms of their host was discovered in the Hippocratic era, but how and why this occurs is enigmatic. Here, we demonstrate that the *Plasmodium chabaudi’s* schedule for asexual replication can be orchestrated by a isoleucine, metabolite provided to the parasite in periodic manner due to the host’s rhythmic intake of food. First, we identify nutrients with daily rhythms in the blood that match the timing of rhythms in both host feeding and the developmental schedule of asexually replicating parasites. We hypothesise that if parasites set their own developmental schedule, they should use a time-of-day cue that is a factor they cannot generate endogenously at any time-of-day, or scavenge in a round-the-clock manner. Our large-scale metabolomics experiment reveals that only one metabolite - the amino acid isoleucine – fits these criteria. Second, further experiments reveal that parasites alter the developmental schedule of asexual stages in response to isoleucine provision and withdrawal in the manner consistent with it acting as a time-cue. Specifically, parasites respond to isoleucine loss by slowing development. This is a parasite strategy rather than the consequences of an imposed constraint, because unlike when parasites are deprived of other essential nutrients, they suffer no apparent costs in the absence of isoleucine. Overall, our data suggest parasites can use the daily rhythmicity of blood-isoleucine concentration to synchronise asexual development with the availability of isoleucine, and potentially other resources, that arrive in the blood in a periodic manner due to the host’s daily feeding-fasting cycle. Identifying both how and why parasites keep time opens avenues for interventions; interfering with the parasite’s time-keeping mechanism may stall replication, increasing the efficacy of drugs and immune responses, and could also prevent parasites from entering dormancy to tolerate drugs.

## Introduction

Circadian rhythms are assumed to have evolved to coordinate organisms’ activities with daily rhythms in the environment (Ouyang et al 1998, Spoelstra et al 2016, Hozer et al 2021). The value of organising sleep/wake cycles according to whether it is light or dark, whether predators or prey are active, *etc*., is clear, but how parasites cope with rhythmicity in the within-host environment has been neglected (Martinez-Bakker & Helm 2015, Reece et al 2017, Westwood et al 2019). Many of the processes underlying interactions between hosts and parasites have a circadian basis, yet why these rhythms exist and what their consequences are for hosts and parasites remain mysterious. For example, parasites are confronted with rhythms in host behaviours and physiologies, including immune responses and metabolism. Thus, some host rhythms offer opportunities for parasites to exploit, whilst other rhythms impose constraints parasites must cope with. Some parasites use their own circadian clocks to control metabolism (Rijo-Ferreira et al 2017), and virulence (Hevia et al 2015), suggesting that host rhythms are a selective (evolutionary) driver of parasite rhythms. Host rhythms have fitness consequences for malaria (*Plasmodium*) parasites (O’Donnell et al 2011, 2013), whose rhythmicity in development during blood-stage replication is aligned with the timing of host feeding-fasting cycles (Hirako et al 2018, Prior et al 2018, O’Donnell et al 2020).

Explaining the timing and synchrony exhibited by malaria parasites during successive cycles of blood-stage replication has been a puzzle since the Hippocratic Era, when the duration of these cycles (24-, 48, or 72-hours, depending on the *Plasmodium spp*.) were used to diagnose malaria infection (Garcia et al 2001, Mideo et al 2013). Blood-stage asexual replication is orchestrated by the intraerythrocytic development cycle (IDC), which is characterised by parasite invasion of red blood cells (RBC), growth, division, and finally bursting of RBC to release the next cohort of asexually replicating parasites. Given that asexual replication is responsible for the disease symptoms of malaria and fuels the production of transmission forms, explaining how and why the vast majority of *Plasmodium* species progress though the IDC in synchrony, and why transitions between these stages occur at particular times of day, may unlock novel interventions and improve the efficacy of existing treatments. However, such endeavours rely on identifying the precise host rhythm(s) that parasites align to and explaining why this matters for their fitness (Prior et al 2020).

Malaria parasites are able to keep time (Rijo-Ferreira et al 2020, Subhudhi et al 2020) and organise their IDC schedule to coordinate with an aspect(s) of host feeding-fasting rhythms (Hirako et al 2018, Prior et al 2018), but not the act of eating itself (Rijo-Ferreira et al 2020), or processes directly controlled by canonical host circadian clocks (O’Donnell et al 2019). Coordination with host feeding-fasting rhythms may allow parasites to couple the nutritional needs of each IDC stage with circadian fluctuations in the concentrations of nutrients in the blood (Prior et al 2020). Whilst parasites are able to scavenge most amino acids from haemoglobin digestion, and can biosynthesise some metabolites themselves, other nutrients are essential and must be taken up from the RBCs or blood plasma. Haemoglobin digestion and biosysnthesis can occur at any time-of-day but essential nutrients that both host and parasite must acquire from the host’s food (including glucose and the amino acid, isoleucine) follow circadian rhythms in the blood (Skene et al 2018). Hosts forage during their active phase and fast during the rest phase, which generally generates peaks in the blood concentrations of metabolites shortly after the onset of feeding and their lowest point (nadir) occurs during the fasting phase.

Nutritional needs vary across IDC stages, with for example, requirements for glycerophospholipids and amino acids to fuel biogenesis, increasing as parasites progress through the IDC, and the release of parasite progeny at the end of the IDC consumes a lot of glucose (Olszewski et al 2009, Déchamps et al 2010, Fig 1A). Whether essential nutrients are available around-the-clock at sufficient concentrations to satisfy the needs of parasites is unknown, but in culture, IDC progression is affected by nutrient limitation. For example, parasites experiencing glucose limitation quickly mount starvation responses (Daily et al 2007) and isoleucine starvation rapidly induces dormancy (Babbitt et al 2012). Moreover, the nadir of blood glucose rhythms in the rest phase can be exacerbated by malaria infection (Hirako et al 2018). Thus, to avoid the problems of temporal resource limitation, we hypothesise that parasites match transitions between IDC stages to rhythmicity in the availability of essential nutrients in the blood, ensuring the most demanding IDC stages (trophozoites and schizonts) coincide with when resources are most abundant. Parasites could synchronise with resource rhythms by using an essential nutrient itself as a time-cue to set the schedule for IDC transitions, and/or allow the host to impose the IDC schedule by, for example, selectively killing IDC stages that are not “on time”. By getting the timing of the IDC schedule right, parasites garner fitness in two ways. They maximise asexual replication which underpins within-host survival (O’Donnell et al 2011, 2013), and benefit from coordinating the production of sexual transmission stages with the time-of-day their vectors forage for blood (Pigeault et al 2018, Schneider et al 2018).

**Fig 1.**
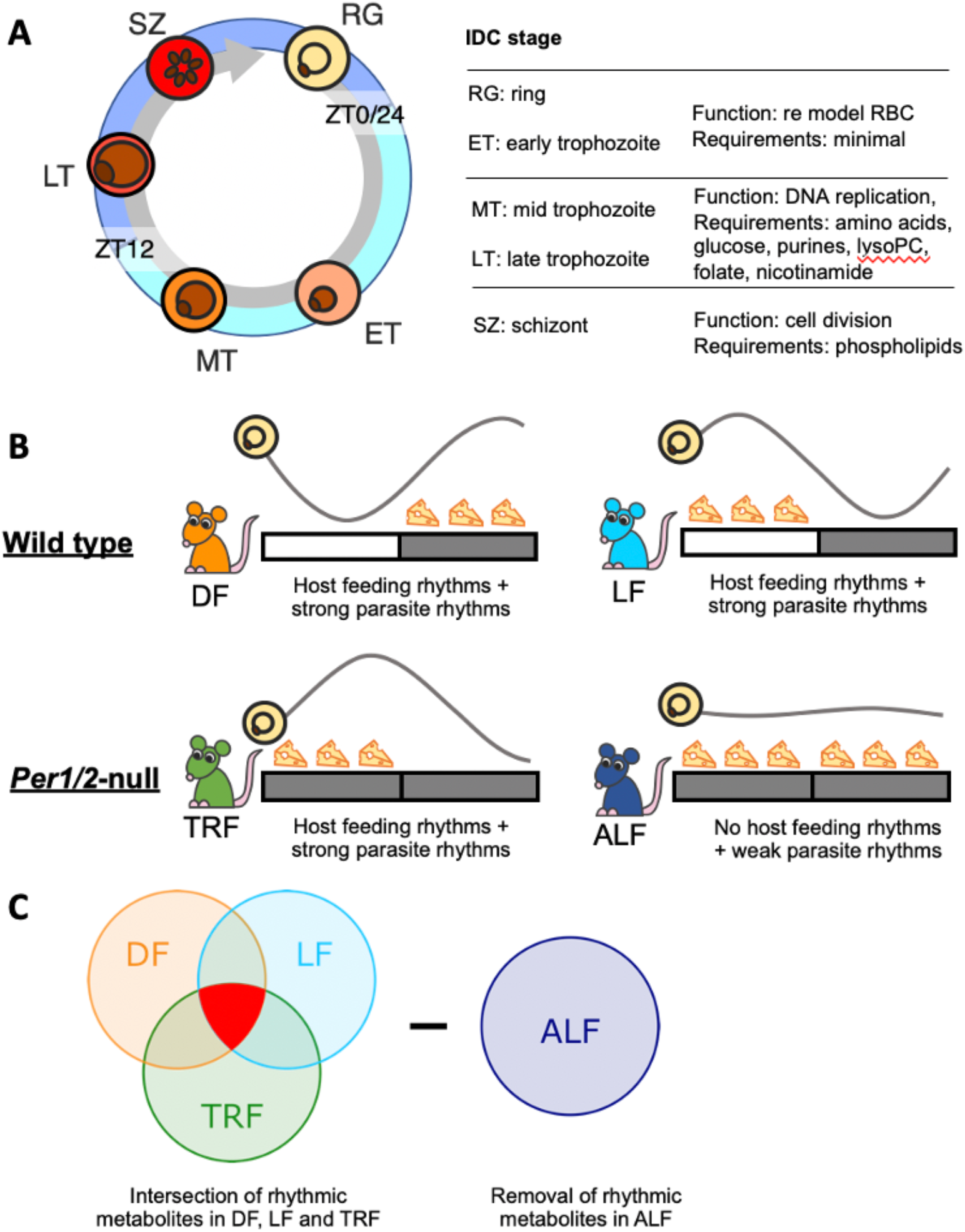
Rhythms in the intraerythrocytic development cycle and host rhythms. **(A)** The role of each IDC stage during cycles of asexual replication and resources known to be essential. RG=ring stage, ET=early stage trophozoite, MT=mid stage trophozoite, LT=late stage trophozoite, SZ=schizont. **(B)**, Experimental design housed wild type mice were in a 12h light: 12h dark regime (indicated by the light-dark bar) with unrestricted access to food for 12 hours either during the night-times (DF, dark feeding) or the day times (LF, light feeding) as indicated by the position of the cheeses. Per1/2-null mice were housed in continuous darkness (DD) and either experienced cycles of 12-hours with food followed by 12 h without access to food (TRF, time restricted feeding,) or given constant access to food (ALF, *ad-libitum* feeding). The parasite IDC is rhythmic in DF, LF and TRF mice but not ALF mice (O’Donnell et al 2019), and so, the IDC rhythm is substantially dampened. **(C)** Metabolites that were significantly rhythmic in DF, LF and TRF mice (highlighted in red), but not in ALF mice, were sought. Treatment groups colour coded throughout: DF=orange, LF=light blue, TRF=green, ALF=dark blue.

We integrate evolutionary ecology and parasitology with chronobiology to, first, undertake a hypothesis-driven screen of several metabolite classes and glucose to identify nutrients with daily fluctuations in the blood of malaria-infected hosts. Second, we determine which metabolites follow rhythms set by the timing of all host feeding-fasting perturbations as well as matching the timing of the IDC schedules in all treatment groups. We find that the IDC schedule cannot be explained by fluctuations in blood glucose concentrations and almost all metabolites, but our screens do reveal a single candidate; the amino acid, isoleucine. Third, we test if the timing of isoleucine provision and withdrawal affects IDC progression in the manner expected if parasites use it as a time-of-day cue to schedule their IDC or if the IDC schedule is imposed by starvation of mis-timed IDC stages. We capitalise on the rodent malaria *P. chabaudi* model system in which *in vivo* experiments exploit ecologically relevant host-parasite interactions and short-term *in vitro* studies allow within-host ecology to be probed in-depth.

## Results

### Metabolites that associate with host feeding rhythms and the IDC schedule

To identify metabolites whose rhythms correspond to the timing of host feeding and the IDC schedule, we compared four groups of malaria infections in mice that were either wild type (WT) C57BL/6J strain or Per1/2-null (Period 1 and Period 2) circadian clock-disrupted mice. Per1/2-null mice have an impaired canonical clock (Transcription Translation Feedback Loop, TTFL) and exhibit no circadian rhythms in physiology or behaviour when kept in constant darkness (Bae et al 2001, Maywood et al 2014; O’Donnell et al 2019). We generated 3 different groups of hosts whose feeding-fasting rhythms differed with respect to the light:dark cycle and whether they had an intact TTFL clock, and a 4^th^ group of hosts which lacked both feeding rhythms and an intact TTFL clock (Fig 1B). All infections were sampled every 2 hours from day 5 post infection (when infections are at a low parasitaemia, ∼10%, to minimise the contribution of parasite molecules to the dataset, Olszewski et al 2009) for 26 hours. We hypothesised that a time-cue/time-dependent resource must vary or have rhythmicity across 24 hours, with a peak concentration that associates with the timing of host feeding-fasting as well as the same IDC stage, across the 3 treatment groups with rhythmic feeding plus a rhythmic IDC, yet be arrhythmic in the 4^th^ group (Fig 1B). Thus, we identified candidate metabolites by intersecting rhythmic metabolites in the treatment groups (Fig 1C).

IDC schedules followed the expected patterns for each treatment group (Fig 2, and O’Donnell et al 2019). Specifically, parasites displayed opposite IDC schedules in dark (i.e. night, DF, n=18) and light (i.e. day, LF, n=17) fed wild type (WT) mice because their feeding-fasting rhythms are inverted (Fig 2, S1 Table). Parasites in Per1/2-null mice (kept in constant darkness) fed during a time-restricted 12 h window (TRF mice, n=17; S2 Table) that coincided with when LF mice were fed, followed the same IDC schedule as parasites in LF mice. Finally, parasites in Per1/2-null mice (kept in constant darkness) with constant access to food (ALF mice, n=16; S2 Table), exhibited dampened IDC rhythms because their hosts have no feeding-fasting rhythm (O’Donnell et al 2019). Across the entire data set, 110 metabolites varied during the 26 hour sampling window (101 in DF, 91 in LF, 50 in TRF and 1 in ALF; S3 Table). That only 1 metabolite (lysoPC a C20:3) exhibited a rhythm in ALF hosts demonstrates that host TTFL clocks and timed feeding are required to generate metabolite rhythms (Fig 3A, Reinke and Asher 2019). Further, that approx. half of the metabolites rhythmic in hosts with TTFL clocks (DF and LF) were also rhythmic in TRF hosts reveals that feeding rhythms alone can maintain metabolic rhythms during infection (Fig 3A). Of all metabolites, only 42 were rhythmic in all 3 groups of infections with both feeding-fasting and IDC rhythms (i.e. the red area in Fig 1C), consisting of 3 acylcarnitines, 11 amino acids, 9 biogenic amines and 19 glycerophospholipids (Fig 3A). Next, we identified 33 metabolites that exhibited a peak phase (timing) that corresponded to the timing of the host’s feeding-fasting rhythm and the parasite’s IDC (Fig 3B; metabolites clustering in the blue and purple areas).

**Fig 2.**
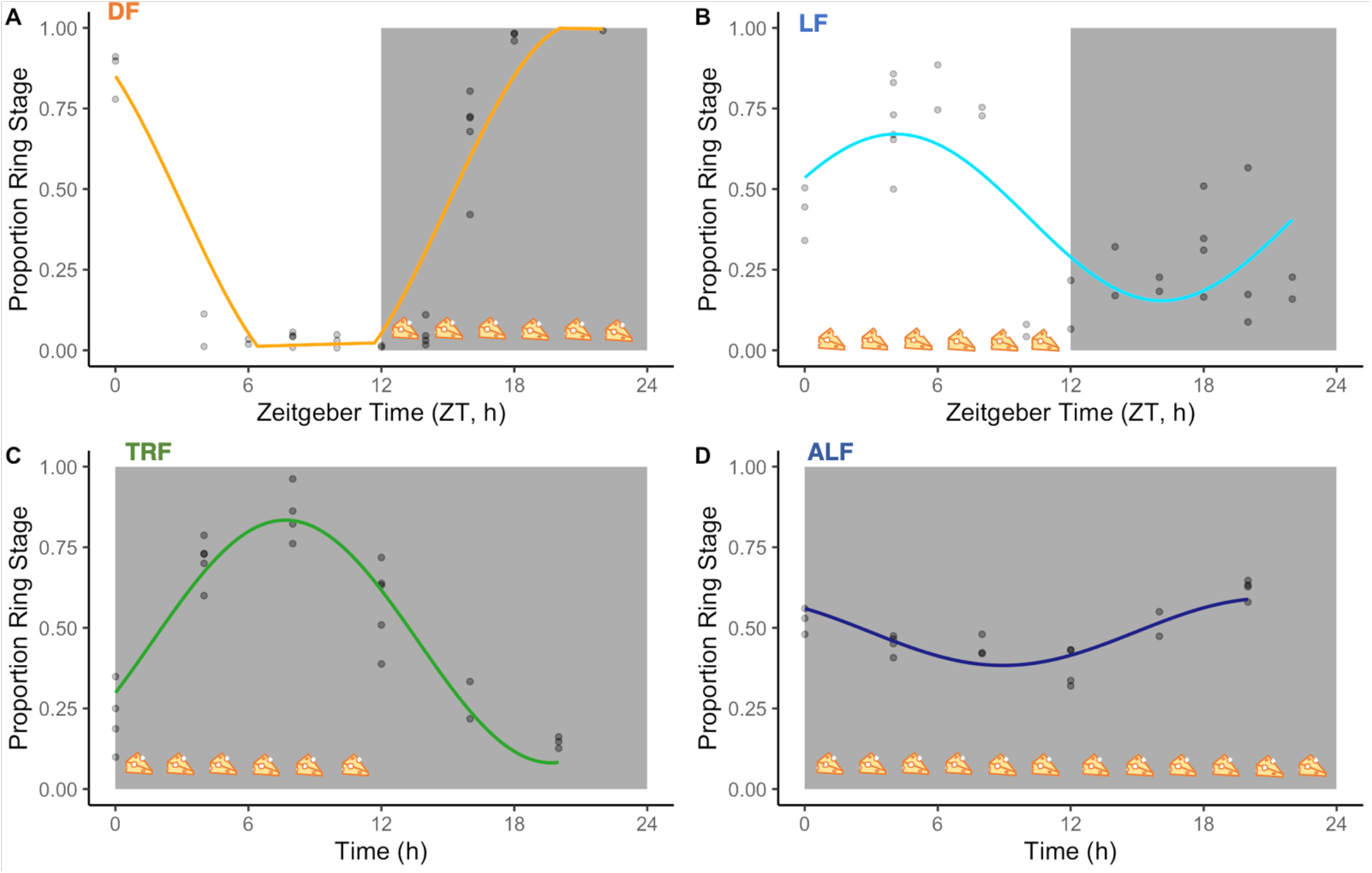
IDC schedule and host feeding rhythms. The proportion of parasites at ring stage as a phase marker, for: **(A)** DF: dark feeding WT mice food access ZT 12-24 (12 h at night) in 12h:12h light:dark, **(B)** LF: light feeding WT mice food access ZT 0-12 (12 h in day) in 12 h:12 h light:dark. **(C)** TRF: time restricted feeding Per1/2-null mice food access 0-12 hours (12 h at the same time (GMT) as LF mice) in constant darkness (DD). **(D)** ALF: ad libitum feeding Per1/2-null mice access in constant darkness (see Fig 1 for Experimental Design and more details). Feeding-fasting rhythms are indicated by cheeses, the white panel denotes lights on (Zeitgeber Time=0-12 h), dark grey panel denotes lights off (Zeitgeber Time=12-24 h for DF and LF, 0-24 h for TRF and ALF). Model fits for each group are plotted on the raw data (n= 2-5 infections per time point). The fitted sine/cosine curve for DF **(A)** is distorted due to a large amplitude which exceeds the bounds possible for proportion data (between 0 and 1) so is truncated for visualisation. The patterns for groups are explained better by sine/cosine curves than by a straight line (see S1 Table),

**Fig 3.**
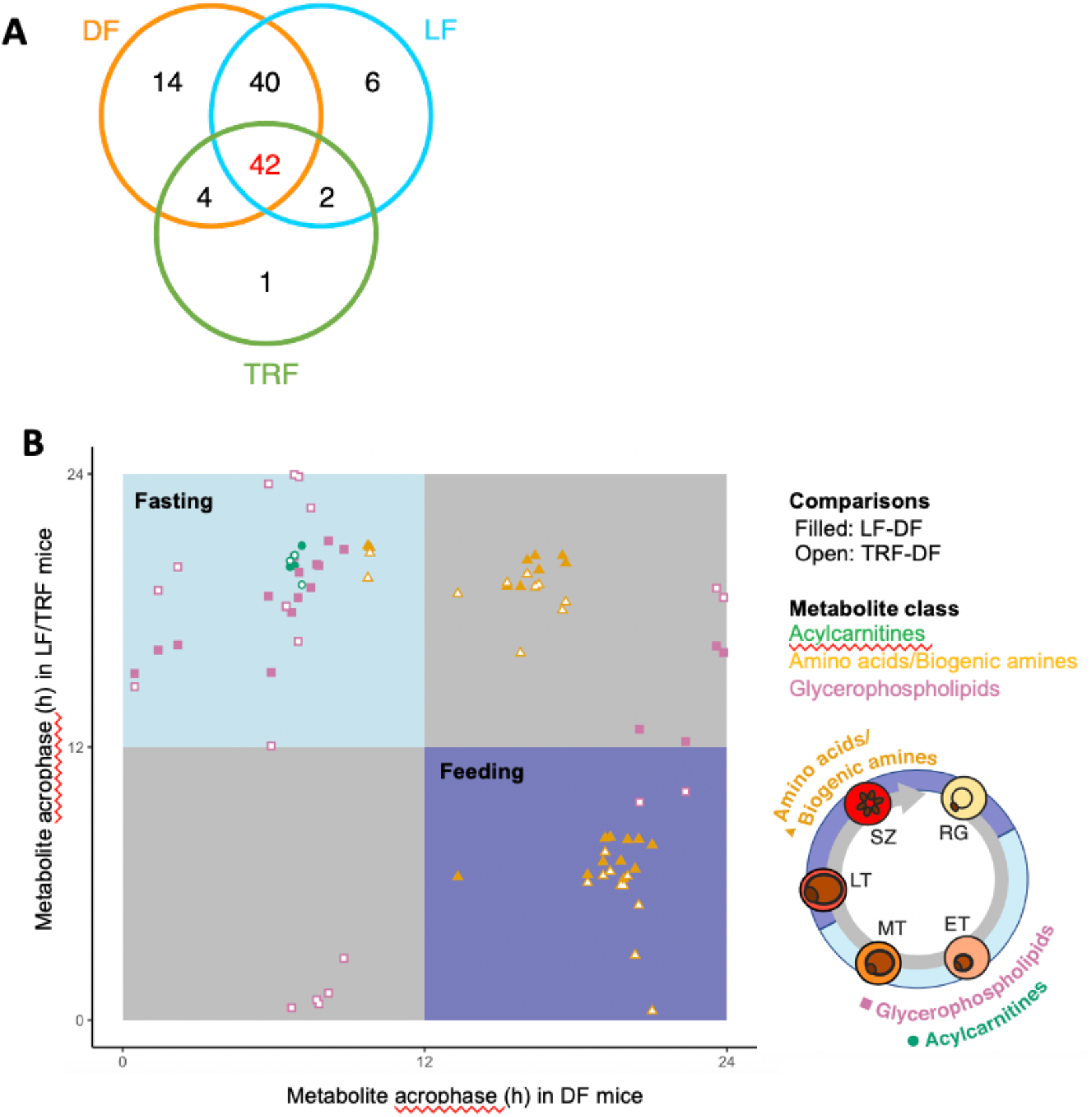
Rhythmic metabolites that associate with IDC timing and host feeding-fasting cycles. **(A)** The numbers of rhythmic intersecting metabolites out of a total of 109. **(B)** Peak timing (phase) concentration of the 42 metabolites that are rhythmic in DF, LF and TRF infections. Top left panel (light blue) denotes peaks during the fasting phase and the bottom right panel (purple) denotes peaks in the feeding phase, yielding 33 candidates linked to the feeding-fasting cycle. Metabolite classes: acylcarnitines=green circles, amino acids/biogenic amines=orange triangles, glycerophospholipids=purple squares. See S5 Table for peak times of each metabolite. Ring stages (RG), late trophozoites (LT) and schizonts (SZ) peak during the hosts feeding period (purple) when some amino acids/biogenic amines peak. Early trophozoites (ET) and mid trophozoites (MT) peak during host fasting (light blue) when glycerophospholipids and acylcarnitines peak.

### Glucose does not associate with the IDC schedule

Previous work suggests that blood glucose rhythms are responsible for the IDC schedule (Hirako et al 2018, Prior et al 2018) but glucose could not be measured by the UPLC/MS-MS metabolomics platform. Therefore, we set up an additional set of DF, LF, TRF and ALF infections (n=5/group), which followed the same feeding-fasting and IDC schedules as infections in the metabolomics screen, Fig 2, S2 Table), to quantify blood-glucose rhythms. Mean blood-glucose concentration differed between the groups, being higher in DF and TRF mice (DF=8.55±0.14 mmol/L, TRF=8.59±0.13 mmol/L) than in LF and ALF mice (LF=7.90±0.14 mmol/L, ALF=7.68±0.11 mmol/L). We found two competitive models including only time of day (Zeitgeber time, ZT/h) or both time-of-day and treatment (DF, LF, TRF and ALF) as main effects, can explain patterns of glucose concentration (Fig 4, S4 Table). Specifically, glucose concentration varied throughout the day in DF mice but much less in LF mice, and glucose was invariant in both groups of Per1/2-null mice (TRF and ALF) (S4 Table). The rhythmic IDC schedule in the DF, LF and TRF groups but the lack of significant rhythmicity in blood glucose in the TRF (and possibly LF) infections demonstrates that glucose does not explain the connection between feeding rhythms and the IDC schedule.

**Fig 4.**
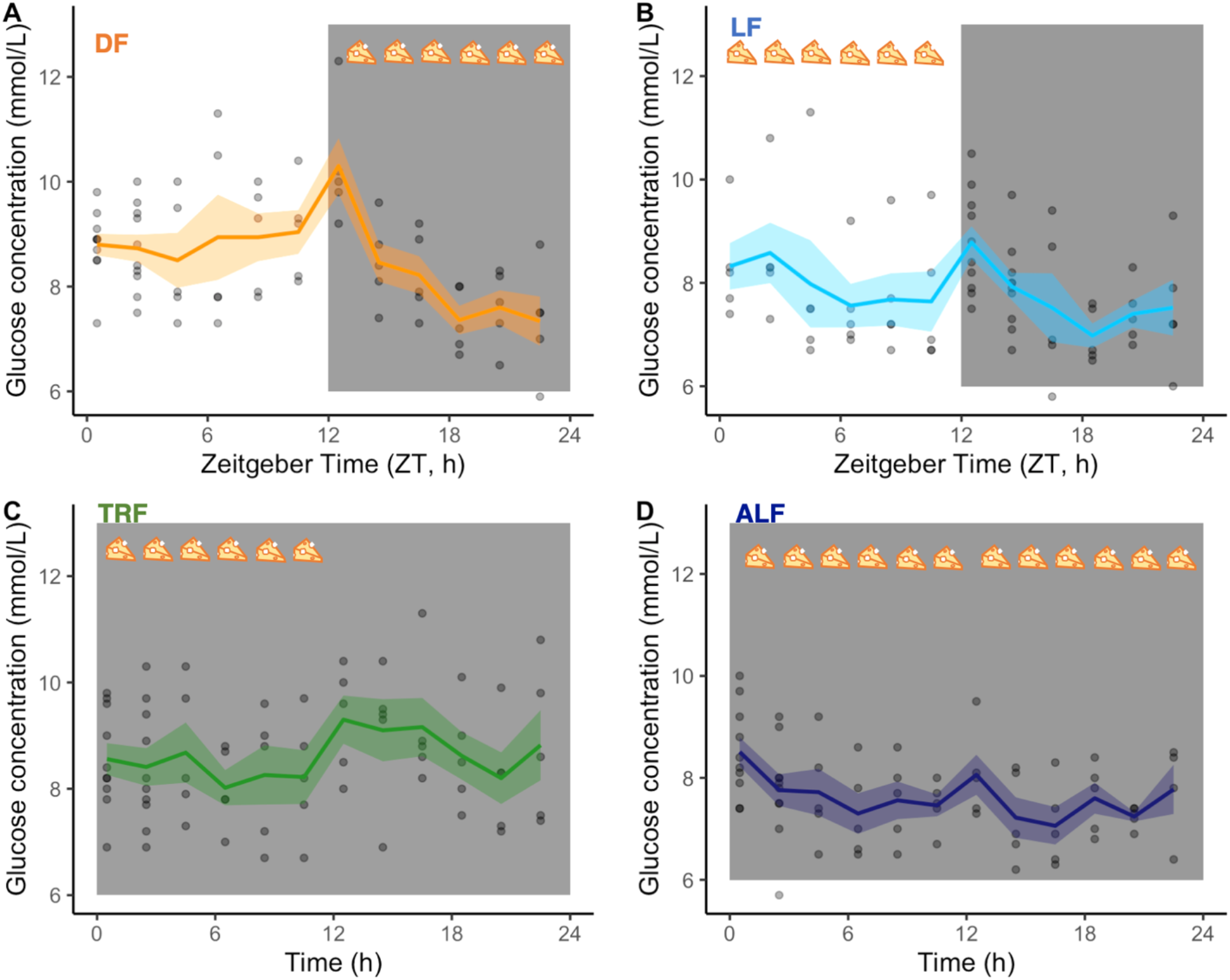
Blood glucose concentrations (mmol/L). **(A)** DF: dark feeding WT mice food access ZT 12-24 (12 h at night) in 12h:12h light:dark, **(B)** LF: light feeding WT mice food access ZT 0-12 (12 h in day) in 12 h:12 h light:dark. **(C)** TRF: time restricted feeding Per1/2-null mice food access 0-12 hours (12 h at the same time (GMT) as LF mice) in constant darkness (DD). **(D)** ALF: ad libitum feeding Per1/2-null mice access in constant darkness (see Fig 1 for Experimental Design and more details). Feeding-fasting rhythms are indicated by cheeses, the white panel denotes lights on (Zeitgeber Time=0-12 h), dark grey panel denotes lights off (Zeitgeber Time=12-24 h for DF and LF, 0-24 h for TRF and ALF). The lines and shading are mean ± SEM at each time point and the dots are the raw data. At time points 0.5 and 2.5 for DF, TRF and ALF, and 12.5 and 14.5 for LF the points are stacked because the time course lasted ∼26 hours and the data are plotted on a 0-24 hour axis.

### Linking metabolites to the IDC schedule

Having ruled out a role for blood glucose, we explored whether the metabolites from our screen could explain why the IDC schedule follows feeding-fasting rhythms. The metabolites whose peak associated with both feeding and late IDC stages, are alanine, asparagine, isoleucine, leucine, methionine, phenylalanine, proline, threonine, valine, methionine-sulfoxide and serotonin. Whereas 20 acylcarnitines and glycerophospholipids (with the exception of 2 biogenic amines) associated with fasting and early IDC stages (ADMA, SDMA, C14.1, C16, C18.1, lysoPCaC16:1, lysoPCaC18:1, lysoPCaC18:2, PCaaC32:1, PCaaC34:4, PCaaC38:3, PCaaC38:4, PCaaC38:5, PCaaC38:6, PCaaC40:4, PCaaC40:5, PCaeC34:3, PCaeC38:0, PCaeC38:3 and PCaeC42:1. S5 Table, S6 Table). Next, we asked if these metabolites could schedule the IDC through the following non-mutually exclusive mechanisms. First, by being sufficiently limiting at a certain time-of-day to enforce a rhythm on the IDC because mis-time IDC stages starve and die. This scenario requires that parasites are unable to overcome resource limitation by synthesising the metabolite itself or scavenging it from a source (such as haemoglobin) that is available around the clock. Second, by acting as a time-of-day cue to which certain IDC stages respond by for example, transitioning to the next stage (called a “just-in-time” strategy) or by acting as a Zeitgeber to entrain an endogenous circadian clock. For a metabolite to be a reliable time-of-day cue/Zeitgeber, it should be something the parasite cannot synthesise/scavenge to avoid the challenge of differentiating between inaccurate endogenous and accurate exogenous time information leading to the risk of mistakenly responding to an endogenous signal. Of the 33 metabolites on the shortlist, only isoleucine fulfils these criteria (Table 1), suggesting that parasites could use isoleucine both as a resource and a time-of-day cue (Fig 5B).

**Table 1.** Potential for acylcarnitines, glycerophospholipids and amino acids/amines candidates that associate with the feeding-fasting cycle to explain the IDC schedule. Requirements for these metabolites is likely to increase as each parasite cell progresses through its IDC.

| Metabolite class & general function | Role in the IDC? | Putative time-cue? |
| --- | --- | --- |
| **Acylcarnitines**<br>• Energy metabolism<br>• Mitochondrial function<br>• Fatty acid transport | No evidence a lack of acylcarnitines affects either the IDC schedule or replication rate.<br>Peak during fasting when parasites are in their least energetically/ metabolically demanding stages. | Very unlikely. |
| **Glycerophospholipids**<br>• Cellular signalling and trafficking<br>• Membrane neogenesis<br>• Haemozoin formation | Phospholipid composition of the host RBC increases during infection, especially oleic acid (18:1) (Déchamps et al 2010).<br>Parasites scavenge fatty acids and also lysoPCs from host plasma, which compete with each other as a source of the acyl components required for lipid synthesis. Lysophosphatidylcholine a C18:1 (lysoPC a 18:1), which contains an oleic acid side chain, is associated with IDC progression.<br>Majority peak during host fasting when parasites are in their least energetically/ metabolically demanding stages. Thus, to be a time-of day cue | Unlikely. Not essential; unlike the liver stage, IDC stages can synthesise fatty acids de novo via type II FA synthase (FASII), (Tarun et al 2009). |
| **Biogenic amines and amino acids:**<br>• Nucleic acid and protein synthesis | Parasites must scavenge several amino acids from the host, including six host-'essential' (isoleucine, leucine, methionine, phenylalanine, threonine and valine) and one host-'non-essential' (alanine) amino acid (Payne and Loomis 2006) that are rhythmic.<br>Parasites can scavenge most amino acids from catabolism of host haemoglobin (Liu et al 2006, Babbitt et al 2012, Martin and Kirk 2007). Isoleucine is the only amino acid absent from human haemoglobin and is one of the least abundant amino acids in rodent haemoglobin (1-3%, Supp Fig 1) yet makes up 9% of both *P. falciparum*'s and *P. chabaudi*'s amino acids (Yadav and Swati 2012).<br>The response of *P. falciparum* to isoleucine withdrawal is not influenced by whether they were previously cultured in a high or low isoleucine concentration environment and results in dormancy that can be broken upon isoleucine replenishment (Babbitt et al 2012).<br>Coinciding with the rise in isoleucine concentration during the feeding window, trophozoites transition to schizonts before bursting and beginning development as ring stages at the end of the feeding window (Fig 5, Fig 7). | Likely. Isoleucine is the only essential amino acid for *P. falciparum*, it cannot be stored, and dormancy has no apparent ill-effects on subsequent development. *P. falciparum* deprived of other amino acids develop as normal (Lui et al 2006) |

**Fig 5.**
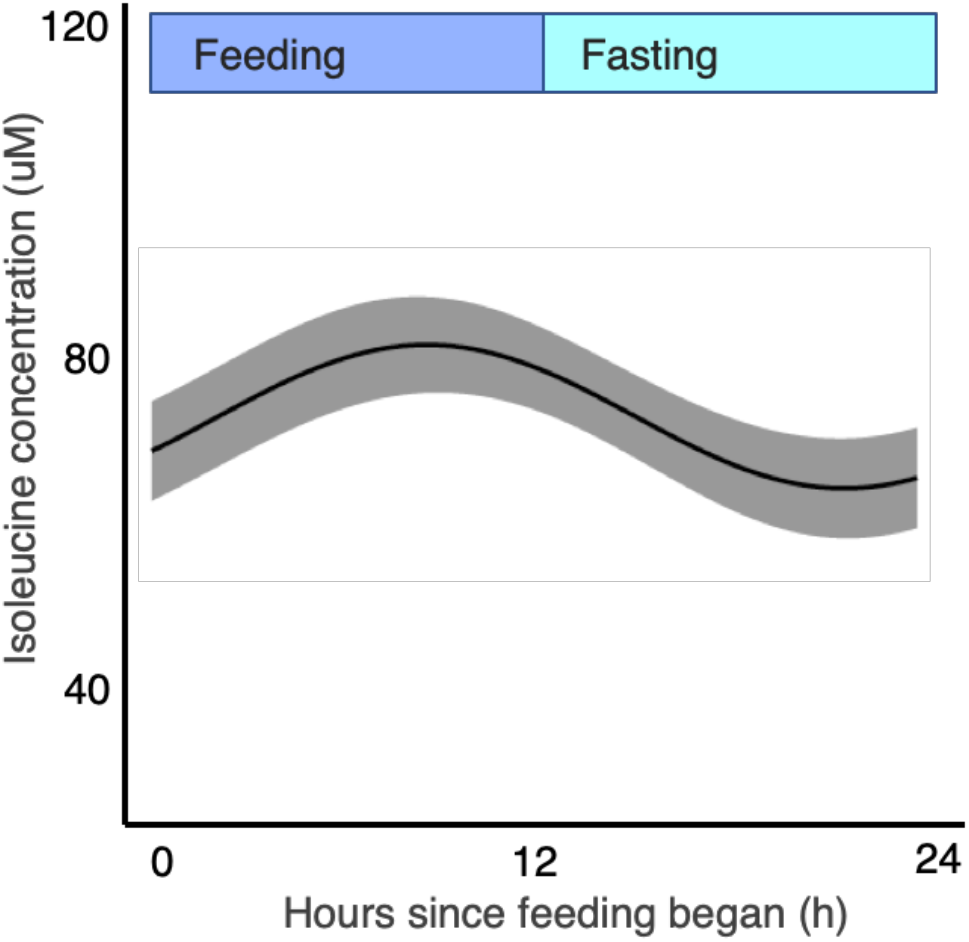
Isoleucine is rhythmic and coincides with the IDC schedule and host feeding-fasting cycles. Model fit (best fit line and 95% prediction interval) for DF, LF and TRF infections combined, from the time since feeding commences, with feeding-fasting windows overlaid.

### Timing and completion of the parasite IDC depends on the availability of isoleucine

That malaria parasites rely on exogenous isoleucine as a both a resource and time-cue to complete the IDC is supported by previous observations across species of *Plasmodium*. First, when the human malaria *P. falciparum* is deprived of isoleucine, parasites immediately and dramatically slow cell cycle progression akin to dormancy (Liu et al 2006) yet are able to recover even after a long periods of starvation (∼4 IDCs) (Babbitt et al 2012). Second, in contrast to isoleucine, if other amino acids are removed from culture media, parasites switch to scavenging them from haemoglobin with only minor effects on IDC completion (Babbitt et al 2012). Third, not only is isoleucine crucial for the development of *P. knowlesi* in culture, parasites only incorporate isoleucine for part of the IDC (until the point that schizogony starts (Polet 1968, Polet 1969, Butcher and Cohen 1971, Sherman 1979)). Fourth, isoleucine is one of few amino acids that exhibits a daily rhythm in the blood of mice and humans. Specifically, isoleucine was inverted with a 12 h shift in simulated day and night shift working humans (a phase difference of 11:49 ± 02:10 h between the day and night shift conditions) and follows the timing of food intake (Skene et al 2018). Therefore, our next experiments tested whether an exogenous supply of isoleucine is capable of scheduling the IDC.

We carried out two experiments in parallel to quantify how *P. chabaudi’s* IDC progression is affected when deprived of isoleucine, and whether the IDC is then completed (defined as the proportion of parasites that reach the schizont stage, S1 Fig) when isoleucine is restored. Isoleucine cannot be perturbed *in vivo* without causing confounding off-target effects on the host or interference by host homeostasis, so we carried these experiments *in vitro* where conditions can be controlled. First, parasites cultured in the absence of isoleucine (n=32 cultures from the blood of 8 mice, which were split equally across both treatments) develop extremely slowly with approx. 3-fold fewer completing the IDC compared to parasites with isoleucine (50 mg/L, which is the same concentration as standard culture media; RPMI 1640) (n=32 cultures) (Fig 6A). The best fitting model contained only “treatment” (parasites cultured with or without isoleucine) as a main effect (ΔAICc=0, S7A Table). The reduction in schizonts in isoleucine-free conditions was not due to a higher death rate because the density of parasites remains constant during the experiment and did not differ between the treatments (Fig 6B) (best fitting model is the null model ΔAICc=0, S7B Table). Further, incorporating either “treatment” or “hours in culture” into the model did not improve the model fit (treatment: ΔAICc=4.16, hours in culture: ΔAICc=5.50, S7B Table).

**Fig 6.**
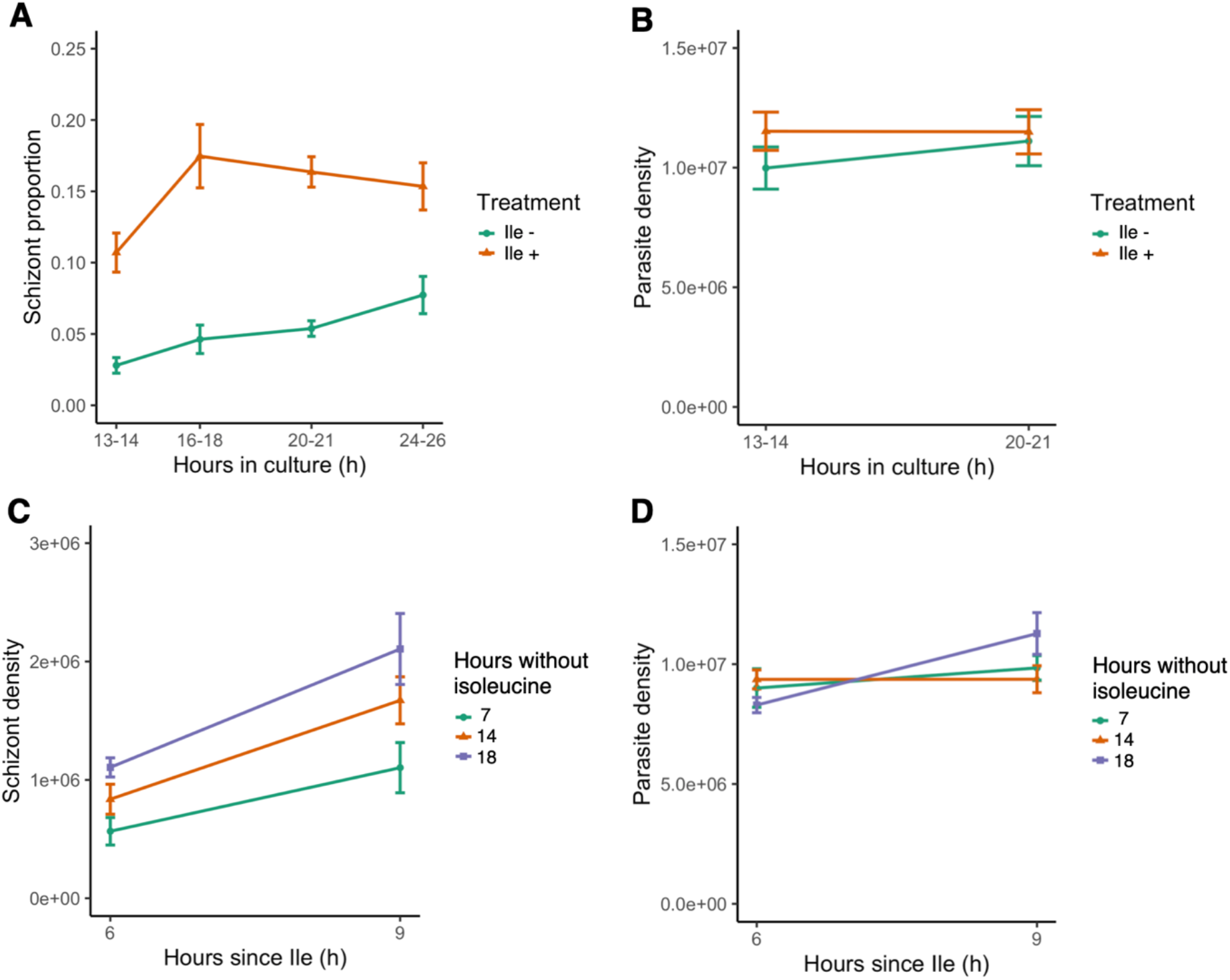
Isoleucine provision and withdrawal drives IDC completion but does affect parasite mortality. **(A)** IDC completion defined as the proportion of parasites that are schizonts, in cultures with (orange triangles, Ile +, 50 mg/L) or without (green circles, Ile -) isoleucine. **(B)** Density of all parasite stages when parasites are cultured with or without isoleucine. Density of **(C)** schizonts and **(D)** all parasite stages, after the addition of isoleucine into cultures following isoleucine deprivation for 7 (green circles), 14 (orange triangles) and 18 hours (purple squares). The proportion of schizonts in the blood seeding the cultures was ∼0.005.

A substantial slowing of IDC progression in the absence of isoleucine is consistent with observations of *P. falciparum* (Liu et al 2006, Babbitt et al 2012, McLean and Jacobs-Lorena 2020) and further supported by our second experiment. This experiment revealed that like for *P. falciparum, P. chabaudi* completes IDC development when isoleucine deprivation ends. Parasites (∼10^7^ per culture) were added to isoleucine-free media and incubated for 7, 14, or 18 hours, after which isoleucine (50 mg/L) was added to their cultures (n=16 cultures per treatment). Parasites completed development when isoleucine became available, regardless of the duration of deprivation (7, 14, or 18 hours), with the best fitting model containing main effects of “treatment” and “hours since isoleucine added” (ΔAICc=0, S7C Table). Importantly, including the interaction did not improve the model fit (ΔAICc=13.65, S7C Table), demonstrating that IDC completion proceeds at the same rate despite different durations of isoleucine starvation. Specifically, the rate of IDC completion in the 6-9 hours following isoleucine replenishment was approximately 50% for all groups (Fig 6C). Again, higher death rates in cultures deprived of isoleucine for the longest time period did not give the appearance of equal IDC rates because the model incorporating “hours since isoleucine” was competitive with the null model (ΔAICc=0.56, Fig 6D, S7D Table), revealing parasites were still as viable after 18 hours of deprivation as those deprived for 7 and 14 hours. Furthermore, cultures deprived the longest achieved the most schizonts (18 hours, mean±SEM: 1.61×10^6^ ±0.20 Fig 6C), while the least schizonts were observed in cultures deprived for the shortest period (7 hours, mean±SEM: 0.83×10^6^ ±0.14 Fig 6C). The variation in the intercepts of Fig 6C (<1% schizonts after 7 hours, 3% after 14 hours and 5% after 18 hours deprivation) is likely explained by the 18 hour deprivation cultures accumulating a higher proportion of schizonts at the time of isoleucine provision simply as a product of developing very slowly during a longer window of deprivation (in line with Lui et al 2006 and Babbitt et al 2012). The behaviour of parasites during isoleucine deprivation and replenishment add further support to isoleucine acting as time-cue as well (Fig 7A) as well as being an essential resource (Fig 7B).

**Fig 7.**
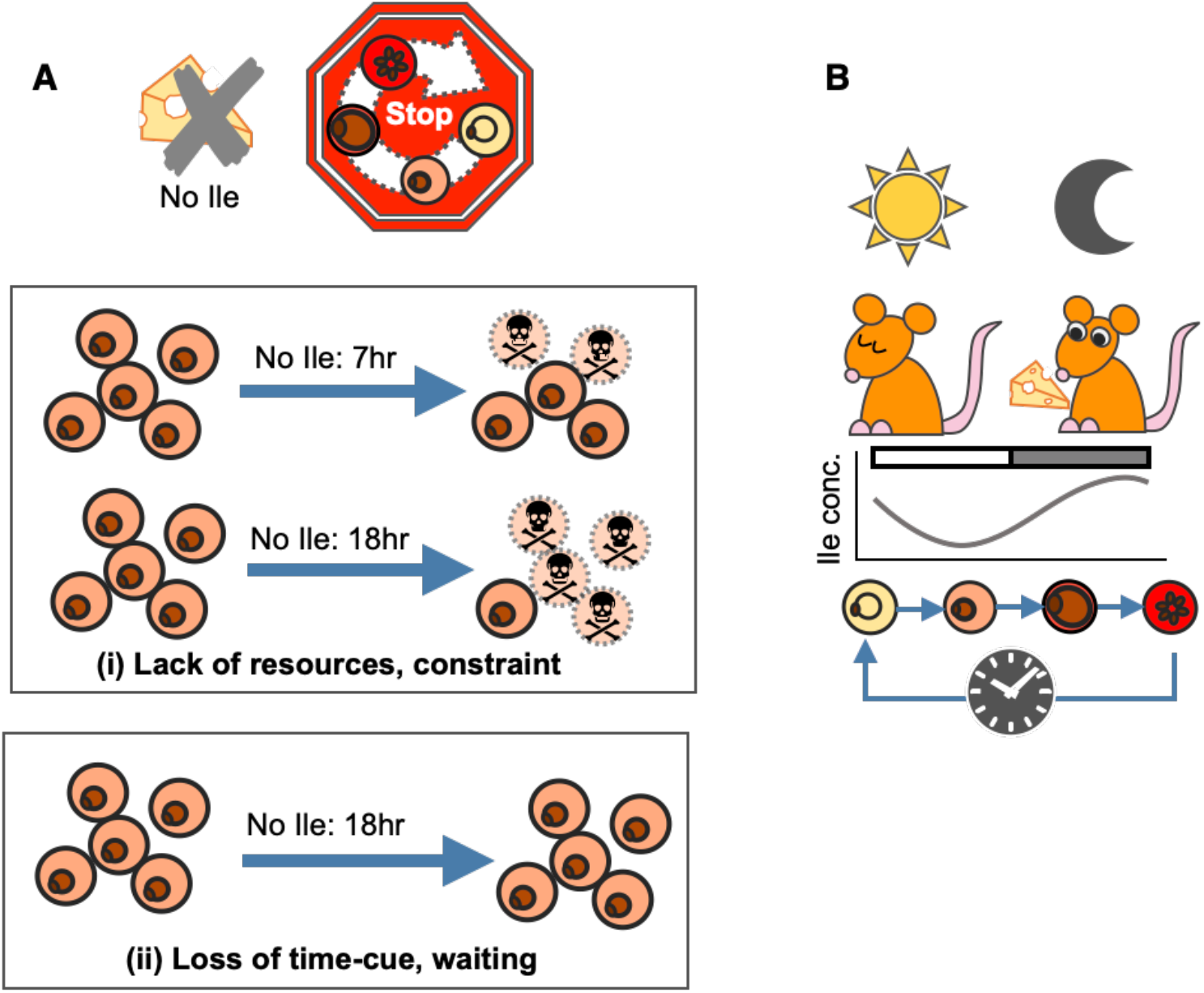
Schematic for isoleucine’s role in the IDC schedule. **(A)** In the absence of isoleucine (Ile), but in the presence of all other essential components of culture media, *P. chabaudi’s* IDC progression stops or continues very slowly, as observed for *P. falciparum* (Lui et al 2006 and Babbitt et al 2012). This observation is consistent with isoleucine being an essential resource and/or a time-cue. This means the time-of-day cue may also be a direct selective agent (i.e. the reason why a cue-response strategy has evolved). In such situations it is very difficult to unravel the consequences of cue loss. Here, the challenge is differentiating to what extent the absence of isoleucine (i) creates a non-permissive environment, imposing a constraint on parasites that simply forces IDC progression to stop, versus (ii) parasites responding in a strategic manner and ‘deciding’ to stop development. Under scenario (i) parasites experience resource limitation, and starvation by definition should have negative consequences. For example, the longer parasites are starved, the more die, or experience increasingly lower or slower rates of recovery upon isoleucine replenishment. Under scenario (ii) there are no such costs because parasites avoid starvation by stopping/slowing development and waiting for a signal that it is time to expect resources to be replenished. That parasite number is not affected by isoleucine withdrawal, even for prolonged periods, and that parasites recover IDC completion at the same rate regardless of the duration of withdrawal, is consistent with scenario (ii) not scenario (i). **(B)** It is now well established that *P. chabaudi’s* IDC schedule is synchronised to host feeding-fasting cycles. Our findings suggest this coordination is achieved by parasites responding to daily rhythmicity in the blood concentration of isoleucine for the purpose of maximsing exploitation of the host’s isoleucine, and potentially other nutrients, they can more easily acquire from the host’s digestion of its food than by biosynthesis or scavenging from haemoglobin.

## Discussion

Our large-scale metabolomics screening experiment (Figs 3-5, Table 1) and follow-up experiments (Fig 6), demonstrate that isoleucine is sufficient to explain how malaria parasites schedule their IDC in line with host feeding-fasting rhythms. A key challenge was differentiating to what extent the absence of isoleucine creates a non-permissive environment, imposing a constraint on parasites that simply forces IDC progression to stop, versus parasites responding in a strategic manner and ‘deciding’ to stop development (Fig 7A). There are several reasons why our findings are not a case of revealing only that parasites cannot develop when resource limited. First, when deprived of other essential resources such as glucose, *P. falciparum* rapidly displays stress responses at the transcriptional level yet when deprived of isoleucine its normal transcriptional pattern persists, albeit very slowly (Babbitt et al 2012), suggesting it is waiting for a signal to proceed (a “gate”). Second, parasites cannot cope with deprivation of other essential resources, for example dying within hours of glucose deprivation (Babbitt et al 2012), yet the IDC restarts with no apparent ill-effects when isoleucine is replenished (Fig 7A). Third, that parasites cannot create isoleucine stores, nor generate it endogenously, and mount rapid transcriptional responses to changes in exogenous isoleucine concentration (Babbitt et al 2012), are all hallmarks of an informative cue and a sensitive response system. Thus, we propose that either parasites are so well adapted to isoleucine starvation they cope just fine without it, unlike their ability to cope with other forms of starvation, or that isoleucine exerts its effects on IDC progression as both a time-cue and resource (Fig 7B). Most studies of isoleucine uptake and use in malaria parasites focus on *P. falciparum*. This parasite uses several channels and receptors (both parasite- and host-derived) to acquire resources from the host. Uninfected human RBC take up both isoleucine and methionine via the saturable L-system (Cobbold et al 2011), which supplies 20% of the necessary isoleucine (Martin and Kirk 2007). When parasitised, there is a 5-fold increase in isoleucine entering RBC which is attributable to new permeability pathways (NPPs) introduced into the RBC membrane by the parasite (Martin and Kirk 2007, McLean and Jacobs-Lorena 2020). NPPs supply 80% of the necessary isoleucine (Martin and Kirk 2007) and are active only in the host membrane at the trophozoite and schizont stages of the IDC, suggesting this influx of isoleucine occurs only at certain times of day (e.g. after host feeding) (Kutner et al 1985). Assuming *P. chabaudi* has analogous mechanisms, we propose that elevated isoleucine is used by the parasite as a marker for a sufficiently nutrient-rich environment to traverse cell cycle checkpoints and complete the IDC (McLean and Jacobs-Lorena 2020, O’Neill et al 2020).

Glucose has previously been suggested as a time-cue or scheduling force for the IDC (Hirako et al 2018, Prior et al 2018). Our data do not support a direct effect of glucose but it may be indirectly involved. Parasites that are glucose limited fail to concentrate isoleucine (Martin and Kirk 2007), likely due to a lack of glycolysis and ATP production needed to operate isoleucine transporters. High concentrations of isoleucine in the blood are also associated with uptake of glucose by tissues, potentially contributing to the hypoglycaemia associated with TNF-stimulation of immune cells during malaria infection (Hirako et al 2018). Additionally, rodent models and humans with obesity and type 2 diabetes like pathologies have elevated levels of isoleucine and dampened glucose rhythms in the blood (Lynch and Adams 2014, Isherwood et al 2017). Thus, if glucose limitation or elevation interferes with the parasite’s ability to acquire time-of-day information from isoleucine, the IDC schedule will be disrupted as observed in diabetic mice (Hirako et al 2018). Connections between isoleucine and glucose might also explain why the parasite protein kinase ‘KIN’ is involved in nutrient sensing (Mancio-Silva et al 2017).

Given that daily variation in the concentration of isoleucine in the blood of infected mice appears modest (55 μM to 80 μM from nadir to peak; Fig 5), we suggest that like *P. falciparum, P. chabaudi* is very sensitive to changes in isoleucine levels. This observation could be interpreted in 2 non-mutually exclusive ways. Perhaps this seemingly damp rhythm means that sufficient isoleucine is available around-the-clock to meet the parasites resource-needs (from the blood and scavenging from the very low level (<3%) of isoleucine in rodent haemoglobin; S1 Fig), so isoleucine acts on the IDC schedule only via a time-cue rather than as a developmental-rate limiting resource. Or, perhaps the isoleucine rhythm becomes exacerbated as infections develop high parasite burdens and hosts become sick. In this case, by aligning the IDC schedule correctly early in infections, parasites minimise the costs of resource limitation later on. In addition to isoleucine itself being essential, its appearance signals a window during which additional factors such vitamins, cofactors, purines, folic acid, pantothenic acid, and glucose that are required for successful replication are also available (Sherman 1979, Müller and Kappes 2007, Müller et al 2010). Furthermore, isoleucine is also reliable cue for the timing of other nutrients that are perhaps easier for parasites to acquire from the host’s digestion of food than from biosynthesis or scavenging from haemoglobin. In particular, that the expression of genes associated with translation are the most commonly disrupted when *P. chabaudi’s* rhythms are perturbed away from the host’s rhythms (Subhudi et al 2020), suggests parasites align the IDC schedule with the resources required for proteins.

Our findings offer a route into identifying the molecular pathways involved in transducing environmental signals to the IDC schedule. To date, the only candidate player identified is SR10, which modulates IDC duration in response to perturbations of host time-of-day (Subudhi et al 2020). Identifying KIN (Mancio-Silva et al 2017), regulated pathways whether they, along with pathways associated with SR10 (Subhudi et al 2020), and PK7/MORC (which is involved in IDC stage transitions; Singh et al 2021), are sensitive to isoleucine might reveal how the parasites’ time-keeping mechanism operates. Whilst by no means conclusive, our results feed the debate about whether malaria parasites possess an endogenous circadian clock. The hallmarks of an endogenous oscillator are (i) temperature compensation, (ii) free running in constant conditions, and (iii) entrainment to a Zeitgeber (Pittendrigh 1960). Recent observations are consistent with (ii) (Subudhi et al 2020, Rijo-Ferreira et al 2020, Smith et al 2020) and our results now allow entrainment to isoleucine to be tested for (iii), as well as free-running in isoleucine-constant conditions (ii), to determine whether malaria parasites possess a clock or use a simpler, reactionary strategy. However, a fast reaction to the withdrawal of isoleucine may not be consistent with a clock. This is because clocks are slow to catch up with a change in the phase (timing) of the Zeitgeber (which is why jet lag occurs), and free run in constant conditions. Thus, if the absence of isoleucine creates free-running conditions, IDC progression should follow the parasites clock and continue as best as possible despite isoleucine limitation, presumably leading to starvation and a loss of number (which we did not observe). Furthermore, most clocks are set by a Zeitgeber (usually light) that is a good proxy for something (e.g. predation risk) that drives selection for a timing strategy. Yet, isoleucine appears to be both cue and selective driver, suggesting a just-in-time reactionary response. This strategy has the advantage of cueing in to the most accurate information about resource availability, and providing a more flexible strategy than a clock when the bulk of host foraging could occur anytime within the broad window of the active phase.

Scheduling development according to the availability of the resources needed to produce progeny intuitively seems like a good strategy to maximise fitness. Yet, the costs/benefits of the IDC schedule demonstrated by parasites may be mediated by parasite density. At low parasite densities (e.g. at the start of infection), resources may be sufficient to support IDC completion of all parasites at any time-of-day, but at intermediate densities, parasites may need to align their IDC needs with rhythms in resource availability. Finally, at very high densities and/or when hosts become sick, resources could be limiting so a synchronous IDC leads to deleterious competition between related parasites. Quantifying whether the costs and benefits of a synchronous IDC vary during infections in line with parasite density and resource availability is complicated by the connection between the IDC schedule and the timing of transmission stages. By aligning the IDC schedule with the feeding-fasting rhythm which is set by light-dark rhythms in the environment, parasites benefit from the knock-on alignment of transmission stage maturation with vector biting activity (O’Donnell et al 2011, Pigeault et al 2018, Schneider et al 2018). Therefore, a key question is to what extent does the IDC schedule contribute to parasite fitness via transmission benefits versus variation during infections in the benefits and costs of aligning asexual replication with rhythmic resources in the blood? More broadly, by integrating evolutionary and mechanistic insight, it may be possible to improve antimalarial treatments. For example, impairing the parasites ability to detect or respond to isoleucine may stall the IDC, reducing virulence and buy time for host immune responses to clear the infection, as well as preventing ring stage dormancy from providing tolerance to antimalarials (Codd et al 20011).

## Acknowledgements

We thank Daan van der Veen, Sam Rund, Petra Schneider, Alejandra Herbert-Mainero and Ronnie Mooney for practical help and discussion.

## Author contributions

Conceptualization, S.R. and K.P; Investigation, K.P., B.M., A.O., M.W., J.H., and A.O.D., Analysis, K.P., D.S., M.B and S.R.; Writing-Original Draft, K.P. and S.R; Writing-Review and Editing, all authors; Supervision, S.R.

## Declaration of Interests

The authors declare no conflicts of interest.

## Methods

### Experimental designs and terminology

The same four perturbations of host and parasite rhythms were used in the metabolomics experiment and the glucose monitoring experiment. The Per1/2-null mice (non-functional proteins Period 1 & 2, backcrossed onto a C57BL/6 background for over 10 generations) were donated by Michael Hastings (MRC Laboratory of Molecular Biology, Cambridge, UK) and generated by David Weaver (UMass Medical School, Massachusetts, USA). Wild type C57BL/6 mice were housed in a 12h light: 12h dark regime (12 hours of light followed by 12 hours of darkness) and the Per1/2-null mice housed in constant darkness (DD) for 2 weeks prior to the start of infections and throughout sampling. We refer to time-of-day using ZT (Zeitgeber Time) for mice housed under entrained conditions (light:dark cycles) and hours when Per1/2-null mice are housed under constant conditions (dark:dark). WT mice either had access to food at night (dark feeding, DF) or in the day (light feeding, LF). Per1/2-null mice either had access to food for 12 hours (time restricted feeding, TRF) or constant access to food (ad libitum feeding, ALF). Every 12 hours, food was added/removed accordingly from the DF, LF and TRF cages and the cages were checked for evidence of hoarding, which was never observed. ALF cages were also disturbed during food removal/provision of the other groups.

#### Sampling

All mice were infected intravenously with *P. chabaudi* DK genotype by intravenous injection of with 1×105 infected RBC at ring stage. Sampling started on day 5 post infection and occurred every 2 hours for both the metabolomics and glucose experiments. We made a thin blood smear each time a mouse was sampled to quantify parasite rhythms by counting ∼100 parasites per blood smear using microscopy. Following (Prior et al 2018, O’Donnell et al 2019 and Rijo-Ferreira et al 2020), we used the proportion of ring stages as a phase marker (an estimate of the timing of parasite development in the blood) of parasite rhythms. We calculated amplitude and time-of-day of peak for each treatment group using sine and cosine curves in a linear model to confirm the IDC schedules for each group as used in O’Donnell et al (2019). All procedures complied with the UK Home Office regulations (Animals Scientific Procedures Act 1986; project 483 licence number 70/8546) and approved by the University of Edinburgh.

#### Metabolomics experiment

We infected 68 eight-week-old female mice: 35 C57BL/6 wild type animals (DF and LF mice) and 33 Per1/2-null TTFL clock-disrupted mice (TRF and ALF) (Supp Table 2). We sampled mice in blocks (A-D), meaning each individual mouse was sampled every 8 hours during the 26-hour sampling window, with 14 time points in total. We did not sample each mouse at each sampling point to minimise the total volume of blood being taken. At each sampling point for each designated host, 20 µl blood was taken from the tail vein to provide 10 µl blood plasma for snap freezing using dry ice.

#### Glucose experiment

We infected 20 eight-week-old male mice: 10 C57BL/6 wild type animals and 10 Per1/2-null circadian clock-disrupted mice (as described above, Supp Table 2). We recorded blood glucose concentration from all mice every 2 hours by taking 1 µl blood using an Accu-Chek Performa Nano glucometer (https://www.accu-chek.co.uk/blood-glucose-meters/performa-nano).

### Targeted metabolomics

We quantified metabolites by analysing plasma samples using the AbsoluteIDQ p180 targeted metabolomics kit (Biocrates Life Sciences AG, Innsbruck, Austria) and a Waters Xevo TQ-S mass spectrometer coupled to an Acquity UPLC system (Waters Corporation, Milford, MA, USA) (Isherwood et al 2017, Skene et al 2018). We prepared the plasma samples (10 µl) according to the manufacturer’s instructions, adding several stable isotope–labelled standards to the samples prior to the derivatization and extraction steps. Using UPLC/MS (ultra performance liquid chromatography/mass spectrometry), we quantified 185 metabolites from 5 different compound classes (acylcarnitines, amino acids, biogenic amines, glycerophospholipids, and sphingolipids). We ran the samples on two 96-well plates, randomised the sample order and ran three levels of quality control (QC) on each plate. We normalised the data between the plates using the results of quality control level 2 (QC2) repeats across the plate (n=4) and between plates (n=2) using Biocrates METIDQ software (QC2 correction). Metabolites were excluded if the CV% of QC2 was > 30% or if all 4 groups contained > 25% of samples that were below the limit of detection, below the lower limit of quantification, or above the limit of quantification or blank out of range. The remaining 134 quantified metabolites comprised of 7 acylcarnitines, 19 amino acids, 15 biogenic amines, 79 glycerophospholipids and 14 sphingolipids (see Supp Fig 3).

### Isoleucine response experiments

To test the effect of the amino acid isoleucine on the IDC we compared parasite developmental progression in cultures with and without isoleucine (50 mg/L), as well as after different durations (7, 14, 18 hours) of isoleucine starvation. We set up N = 112 cultures (from 8 mice) so that for each time point within each treatment, an independent culture was sampled, avoiding any bias associated with repeat-sampling individual cultures. We used eight-week-old wild type female mice, MF1 strain, housed in a 12h:12h light:dark regime before and during infection. We infected mice intraperitoneally with 1×106 *P. chabaudi* genotype DK infected red blood cells at ring stage and terminally bled them on day 6 post infection, when the parasitaemia was around 15%. Mice were bled at ZT4, when parasites were late rings/ early trophozoites (see Prior et al 2018). Approximately 1 ml of blood was collected from each mouse which was then split equally across cultures in all 5 treatment groups.

#### Culturing

We washed infected blood twice with buffered RPMI containing no amino acids (following Spence et al 2011, see Supp Mat “Parasite culture protocol” for more details), before being reconstituted in the RPMI medium corresponding to each treatment (containing isoleucine, or not). We cultured parasites in 96-well round bottom plates with total culture volumes of 200-250 µl, at ∼3% haematocrit and kept the culture plates inside a gas chamber which was gassed upon closing with 88% nitrogen 7% carbon dioxide and 5% oxygen, and then placed inside a 37°C incubator. The culture medium was custom made RPMI from Cell Culture Technologies, Switzerland (http://www.cellculture.com) (see Supp Mat “Parasite culture protocol”).

#### Sampling

We sampled parasites in the first experiment (comparing IDC completion in isoleucine rich versus isoleucine free media) at 13-14, 16-18, 20-21, 24-26 and 27 hours after culture initiation. We also sampled parasites that had been isoleucine deprived for 7, 14 or 18 hours at 6 and 9 hrs after isoleucine addition. Samples consisted of a thin blood smear from each culture fixed with methanol and Giemsa stained. We measured the proportion of parasites in the schizont stage (as an indicator of parasites having completed their IDC) by counting ∼300 parasites per blood smear.

### Statistical analysis

We defined daily rhythmicity in the concentrations of the 134 metabolites detected by the UPLC/MS-MS platform as the detection of rhythmicity in at least 2 of the following circadian analysis programmes: ECHO (https://github.com/delosh653/ECHO), CircWave (https://www.euclock.org/results/item/circ-wave.html) and JTK_Cycle (https://openwetware.org/wiki/HughesLab:JTK_Cycle). For metabolites not found to be rhythmic by at least 2 of the circadian analysis programmes, we also carried out Analysis of Variance to identify metabolites that varied across the day but without a detectable 24 h rhythm (Supp Table 3 for breakdown of which candidate metabolites were rhythmic in each programme). To calculate the acrophase (timing of peak) we used linear mixed-effects regression models containing sine and cosine terms on all metabolites that varied across the day. Metabolites whose acrophase fell in the same 12h feeding or fasting window (ZT0-12 or ZT12-24) in LF and TRF infections but fell in the opposite window for the DF group were shortlisted as potential candidates to connect host feeding rhythms and the IDC schedule. We compared schizont proportion and the density of combined IDC stages using linear regression models. To avoid overfitting due to small sample sizes, “Akaike information criterion-corrected” (AICc) values were calculated for each model, and a change in 2 AICc (ΔAICc = 2) was chosen to select the most parsimonious model (Bewer et al 2016).

## Supplemental material

### Supplementary Materials and Methods

**Parasite culture protocol**. Modified from Spence et al (2011).

**Equipment**

- Water bath (37°C) (Nickel Electro NE3-28DT)
- Heated centrifuge (37°C) (5810R, Eppendorf, Germany)
- Heat block (37°C) (cat. no. N2400-4020, Star Lab, UK)
- Needles and syringes (27 G × 0.5 inch; BD Microlance 3; BD, cat. no. 300635, GP Supplies, UK)
- 15 ml Falcon tubes (cat. no. CLS430055, Sigma-Aldrich, UK)
- Microscope slides (Menzel-Gläser 8037/1, Thermo Scientific, UK)
- Flow cytometer (Z2 Coulter Counter, Beckman Coulter, US)
- Millipore filter (Millex-GP 33 mm PES .22 um Sterile, Merck, US)
- Incubator (37°C) (Panasonic Programmable Cooled Incubator, MIR-154, PHCbi, Japan)
- Cell culture plates (cat. no. SIAL0799, Sigma-Aldrich, UK)
- Compressed gas mixture: 5% O_2_, 7% CO_2_, 88% N_2_ (BOC, UK)
- Incubator chamber

**Reagents**

- Heparin (cat. no. H3393, Sigma-Aldrich, UK)
- 10% Giemsa’s stain (cat. no. 48900-500ML-F, Sigma-Aldrich, UK) in 1× Giemsa’s phosphate buffer (cat. no. P4417, Sigma-Aldrich, UK)
- Custom-made RPMI 1640: medium kit “basic RPMI 1640” (without amino acids but including glucose, salts and vitamins) (www.cellculture.com), tissue culture water, sodium hydrogen carbonate.
- Ile-medium: basic RPMI 1640 plus glycine (10 mg/L), L-arginine (200), L-asparagine (50), L-aspartic acid (20), L-cystine 2HCl (65), L-glutamic acid (20), L-glutamine (300 - this is added to the basic RPMI), L-histidine (15), L-hydroxyproline (20), L-leucine (50), L-lysine hydrochloride (40), L-methionine (15), L-phenylalanine (15), L-proline (20), L-serine (30), L-threonine (20), L-tryptophan (5), L-tyrosine disodium salt dihydrate (29)), L-valine (20).
- Ile+ medium: Ile-medium plus L-isoleucine (50 mg/L)
- Buffered RPMI: basic RPMI 1640, 2 µM L-glutamine (cat. no. 25030081, Gibco, UK) and 6 mM HEPES (sterile) (cat. no. 15630080, Gibco, UK).
- Complete RPMI: basic RPMI 1640, 2 mM L-glutamine, 6 mM HEPES, 0.5 mM sodium pyruvate (cat. no. 11360070, Gibco, UK), 50 µM 2-Mercaptoethanol (cat. no. 21985023, Gibco, UK), 10 µl gentamicin (cat. no. 15710049, Gibco, UK), 10% Albumax (sterile) (cat. no. 11021037, Gibco, UK).

**Procedure**

1. Inject donor mice intraperitoneally with at least 10^6^ infected RBC, monitoring the infections daily on Giemsa-stained thin blood films.
2. Around day 6 post infection (or when parasitaemia is around 10-15%), heart bleed mice when parasites are at early trophozoite stage ensuring to use heparin to prevent clotting (around 50-100ul) keeping all equipment at 37°C (tubes, syringes).
3. Collect blood from each mouse into 15 ml Falcon tubes. Measure the total volume of collected blood and for every 1 ml of blood add 10 ml prewarmed buffered RPMI and place in 37°C water bath. Wash blood twice by centrifuging at 2200 rpm/1770 G for 5 mins at 37°C in buffered Ile-medium, split blood between treatment groups, then wash for a second time and resuspend in the correct medium (Ile+ or Ile-).
4. Culture parasites at 2-5% haematocrit ((50×2) × **x** ml blood, assuming hematocrit is 50% in whole blood and **x** is the total amount of blood collected) in Ile+ or Ile-medium, and add 250 ul of the final culture concentration to each well on a 96-well round bottom plate.
5. Stack 96-well plates inside incubator chamber, gas, then place inside 37°C incubator.
6. Sample each well as required using a 10 ul pipette by gently scraping the bottom layer of RBCs and make a thin blood smear. Re-gas the chamber before placing the 96-well plates back in the incubator.

## Supplementary Figures

**Supp Fig 1.**
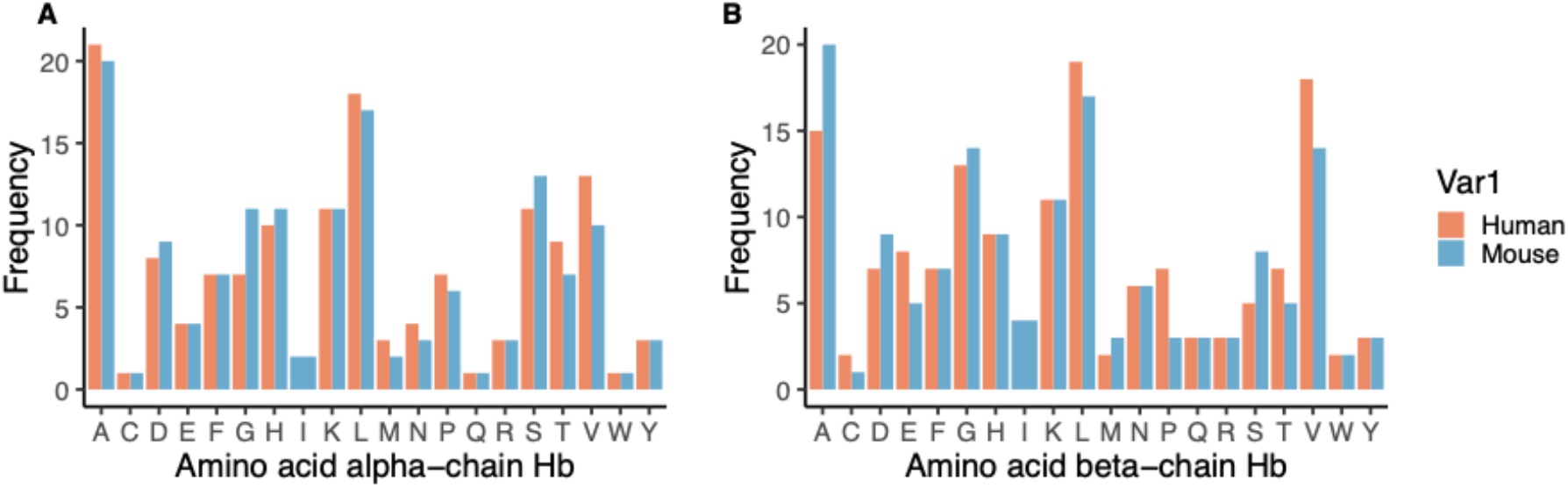
Frequency of amino acids in alpha (A) and beta (B) chains of human (orange) and mouse (blue) haemoglobin. Amino acid codes: A - alanine, C - cysteine, D - aspartic acid, E - glutamic acid, F - phenylalanine, G - glycine, H - histidine, I - isoleucine, K - lysine, L - leucine, M - methionine, N - asparagine, P - proline, Q - glutamine, R - arginine, S - serine, T - threonine, V - valine, W - tryptophan, Y - tyrosine.

**Supp Fig 2.**
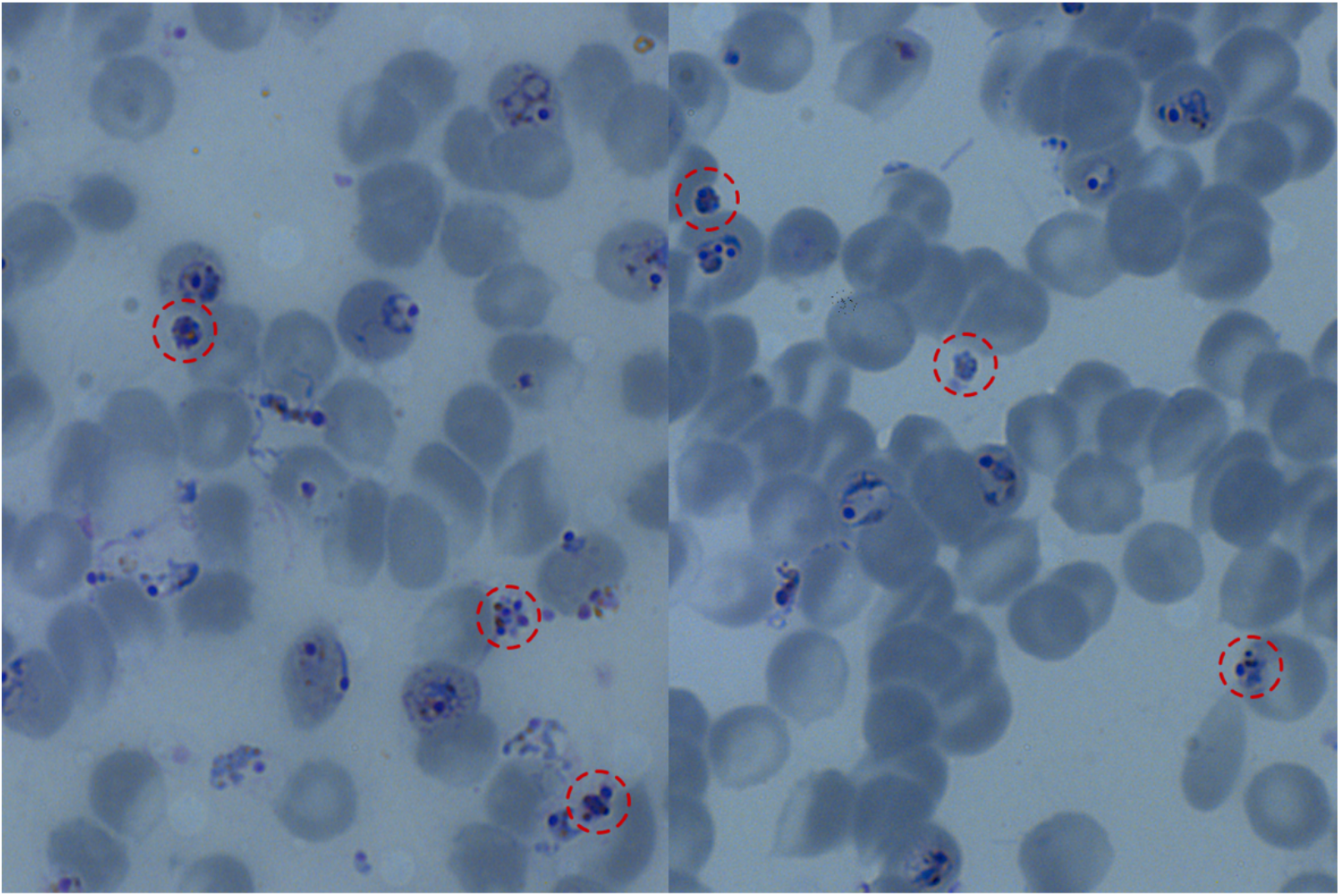
Photo of infected RBCs in *P. chabaudi* culture, schizonts circled in red. Using Giemsa RBCs stain grey and parasite organelles and nuclei stain blue-purple. Schizonts inside and outside of RBCs were counted (assumed to be viable) as RBC membranes are delicate and are easily disrupted during thin smear production. Parasites spent ∼30 h in culture and sampled every 3-4 hours after 13 hours of culture, with each well from the corresponding mouse only being sampled once (n=8 per time point).

**Supp Fig 3.**
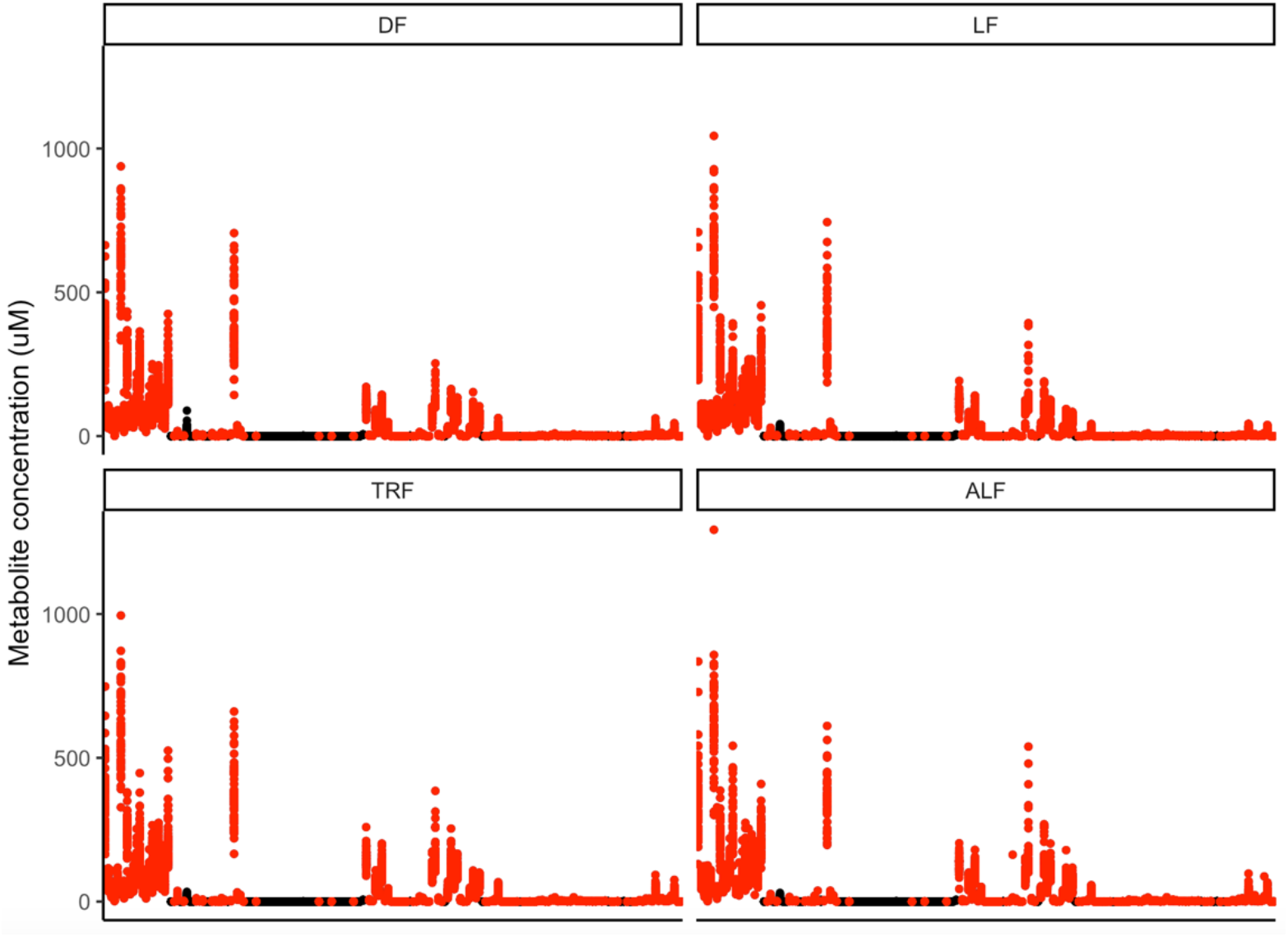
Concentration of all metabolites at all time points during the time series, those included and excluded from the analysis. DF=dark fed wild type mice, LF=light fed wild type mice, TRF=time restricted fed Per1/2-null mice, ALF=ad libitum fed Per1/2-null mice. Each x-axis mark is a different metabolite with concentration (µM) on the y-axis. In black are metabolites excluded since they failed set LC/MS assay criteria and those metabolites taken forward into the analysis are red.

## Supplementary Tables

**Supp Table 1.**
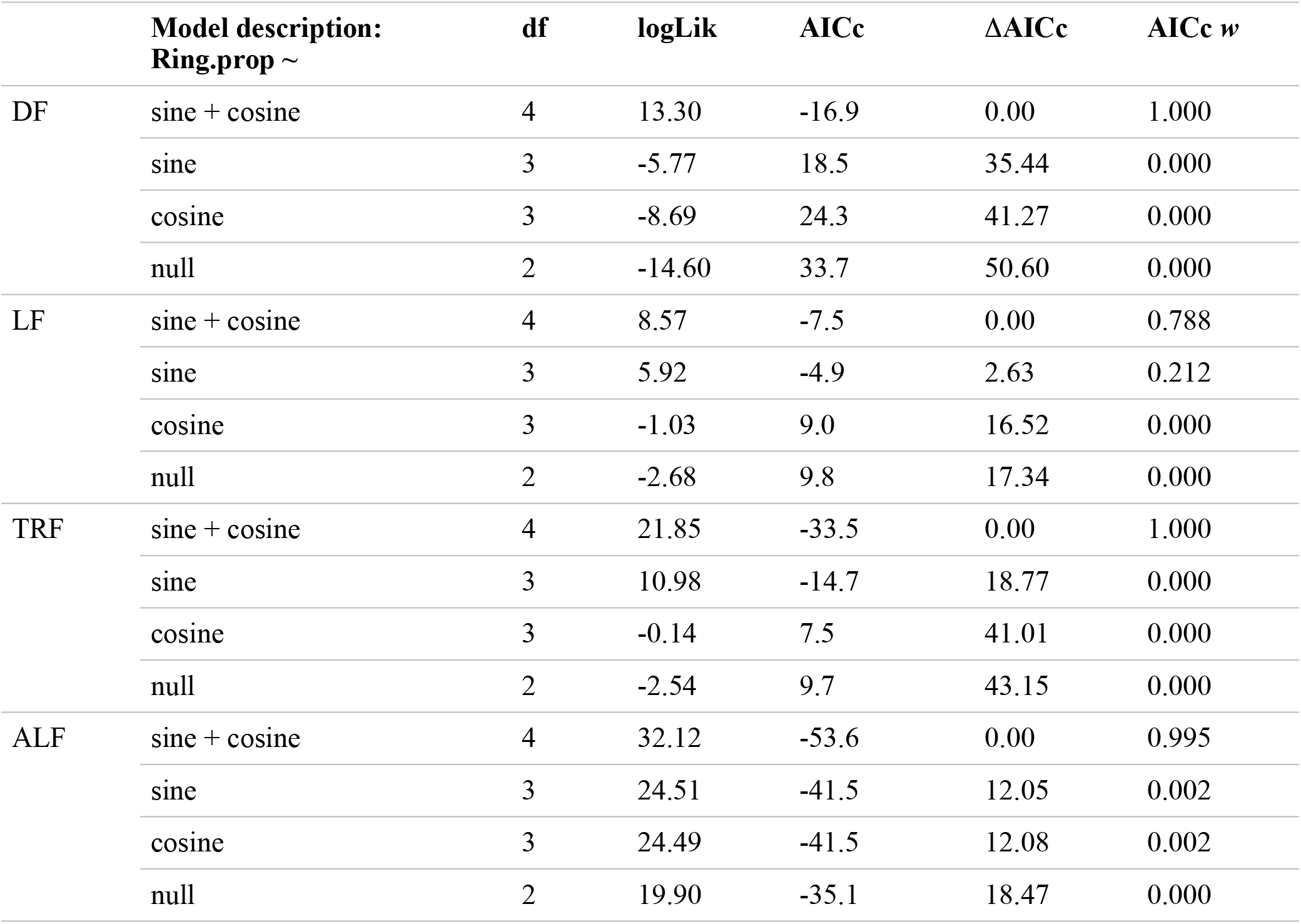
Degrees of Freedom (df), log-Likelihood (logLik), AICc, ΔAICc (AICc_i_ – AICc_min_) and AICc w (AICc weight) for each linear model in the parasite stage proportion analysis ordered in descending fit (best-fitting model at the top). The response variable for each model is proportion of ring stages (Ring.prop), with “sine” and “cosine” terms being the sine or cosine function of (2π × time of day)/24 with a fixed 24h period fitted for each treatment group (DF, LF, TRF or ALF). AICc is a form of the Akaike Information Criteria corrected for smaller sample sizes to address potential overfitting, used for model selection. Corresponds with Fig 2.

**Supp Table 2.** Mouse numbers for experimental treatment groups. A) Mice in the metabolomics experiment were sampled in blocks (A-D) with 4/5 mice per block sampled every 8 hours. Totals for each treatment group: DF=18, LF=17, TRF=17, ALF=16. B) Mice in the glucose experiment were sampled every 2 hours. Totals for each treatment group: DF=5, LF=5, TRF=5, ALF=5. For each experiment repeated measures from mice were controlled for during the analysis. Relates to Figs 1, 2, 4.

| Time (ZT/<br>hour) | 0 | 2 | 4 | 6 | 8 | 10 | 12 | 14 | 16 | 18 | 20 | 22 | 24/0 | 2 |
| --- | --- | --- | --- | --- | --- | --- | --- | --- | --- | --- | --- | --- | --- | --- |
| **A Metabolomics experiment** |  |  |  |  |  |  |  |  |  |  |  |  |  |  |
| *Block* | *A* | *B* | *C* | *D* | *A* | *B* | *C* | *D* | *A* | *B* | *C* | *D* | *A* | *B* |
| DF | 5 | 5 | 4 | 4 | 5 | 5 | 4 | 4 | 5 | 5 | 4 | 4 | 5 | 5 |
| LF | 5 | 4 | 4 | 4 | 5 | 4 | 4 | 4 | 5 | 4 | 4 | 4 | 5 | 4 |
| TRF | 5 | 4 | 4 | 4 | 5 | 4 | 4 | 4 | 5 | 4 | 4 | 4 | 5 | 4 |
| ALF | 4 | 4 | 4 | 4 | 4 | 4 | 4 | 4 | 4 | 4 | 4 | 4 | 4 | 4 |
| **B Glucose experiment** |  |  |  |  |  |  |  |  |  |  |  |  |  |  |
| DF | 5 | 5 | 5 | 5 | 5 | 5 | 5 | 5 | 5 | 5 | 5 | 5 | 5 | 5 |
| LF | 5 | 5 | 5 | 5 | 5 | 5 | 5 | 5 | 5 | 5 | 5 | 5 | 5 | 5 |
| TRF | 5 | 5 | 5 | 5 | 5 | 5 | 5 | 5 | 5 | 5 | 5 | 5 | 5 | 5 |
| ALF | 5 | 5 | 5 | 5 | 5 | 5 | 5 | 5 | 5 | 5 | 5 | 5 | 5 | 5 |

**Supp Table 3.** Metabolite numbers that significantly fluctuate every 24h in the mouse blood for three methods. **A)** Data for all metabolites were run through three circadian programmes to find those following a 24h rhythm using ECHO (Benjamini-Hochberg adjusted p value of 0.05), CircWave (standard p value of 0.05) and JTK_Cycle (BH adjusted p value of 0.05). Metabolites that are excluded using the Surrey LC/MS assay criteria were removed from the analysis (Supp Fig 3). **B)** Rhythmic metabolites in each programme (ECHO, CircWave and JTK) were intersected, with a metabolite counted as rhythmic if it is significantly rhythmic in at least two programmes (ECHO=CW, ECHO=JTK, CW=JTK). **C)** The metabolites not fulfilling these criteria were analysed using ANOVA (including time-of-day as a factor), with BH adjusted p values at the 5% level. Metabolites from both methods (circadian programmes and ANOVA) were then combined to perform a final intersection between DF, LF and TRF to find common metabolites. Metabolites rhythmic in ALF mice were then removed. Relates to Fig. 3.

| **A** | **ECHO** |  | **CircWave (CW)** |  | **JTK** |  |
| --- | --- | --- | --- | --- | --- | --- |
|  | Rhythmic with 24h period | Not-rhythmic with 24h period | Rhythmic with 24h period | Not-rhythmic with 24h period | Rhythmic with 24h period | Not-rhythmic with 24h period |
| DF | 75 | 59 | 67 | 67 | 47 | 87 |
| LF | 106 | 28 | 83 | 51 | 70 | 64 |
| TRF | 55 | 77 | 41 | 93 | 23 | 111 |
| ALF | 2 | 127 | 6 | 128 | 0 | 134 |

| **B** | ECHO==CW | ECHO==JTK | CW==JTK | Unique |
| --- | --- | --- | --- | --- |
| DF | 62 | 47 | 46 | 63 |
| LF | 79 | 68 | 63 | 88 |
| TRF | 37 | 23 | 21 | 39 |
| ALF | 1 | 0 | 0 | 1 |

| **C** | Not-rhythmic with 24h period | Time-of-day effect | No time-of-day effect | Time-of-day effect + rhythmic with 24h period | Total candidates |
| --- | --- | --- | --- | --- | --- |
| DF | 71 | 38 | 33 | 101 | 42 |
| LF | 46 | 3 | 43 | 91 |  |
| TRF | 95 | 11 | 84 | 50 |  |
| ALF | 133 | 0 | 133 | 1 |  |

**Supp Table 4.**
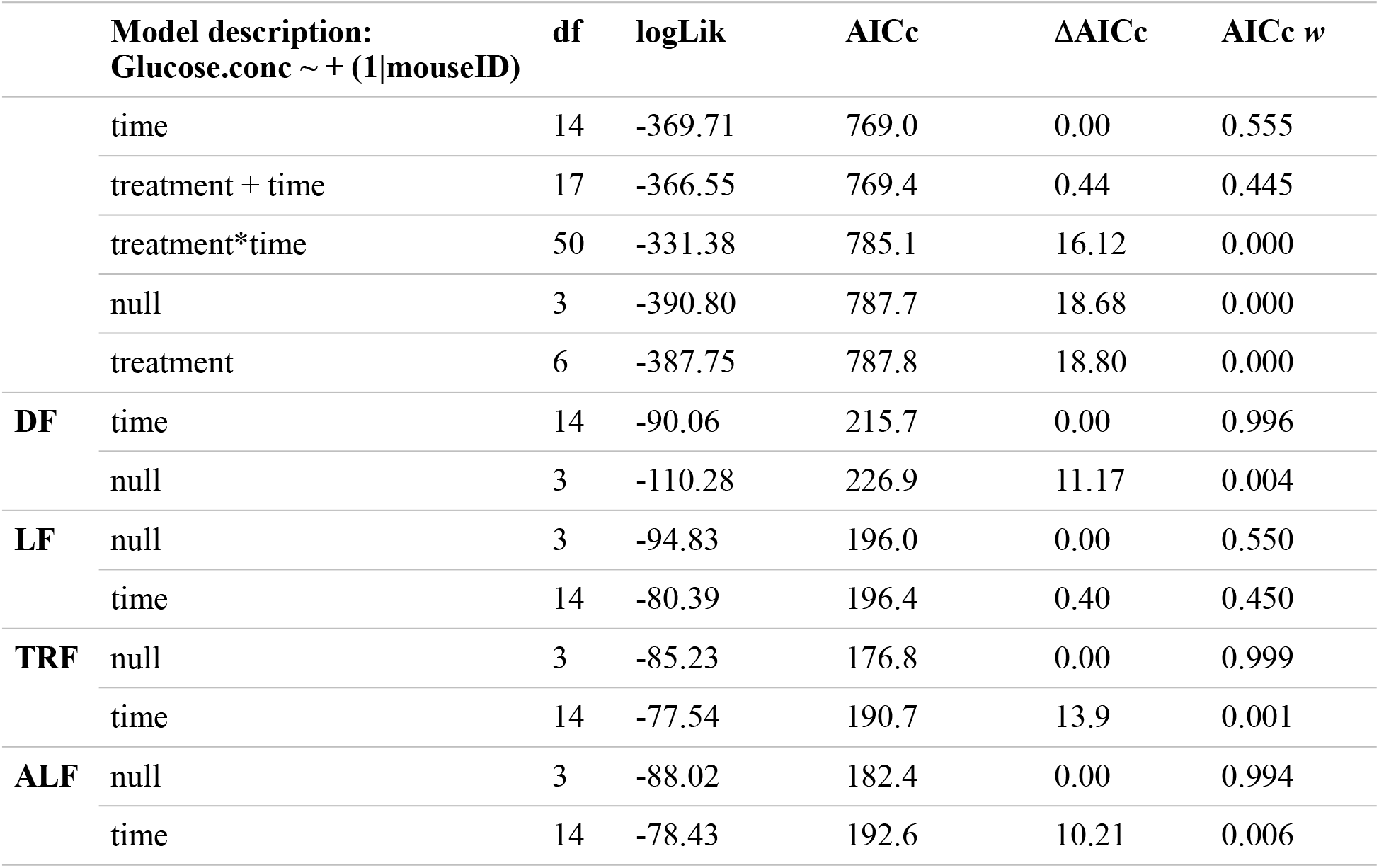
Degrees of Freedom (df), log-Likelihood (logLik), AICc, ΔAICc (AICc_i_ – AICc_min_) and AICc w (AICc weight) for each linear model in the glucose concentration analysis ordered in descending fit (best-fitting model at the top). The response variable for each model is glucose concentration (Glucose.conc) and the random effect is “mouseID”. “Treatment” refers to the treatment group (DF, LF, TRF or ALF) and “time” refers to the time-of day (ZT 0-24h or 0-24h) which was fitted as a factor. Relates to Fig 4.

**Supp Table 5.**
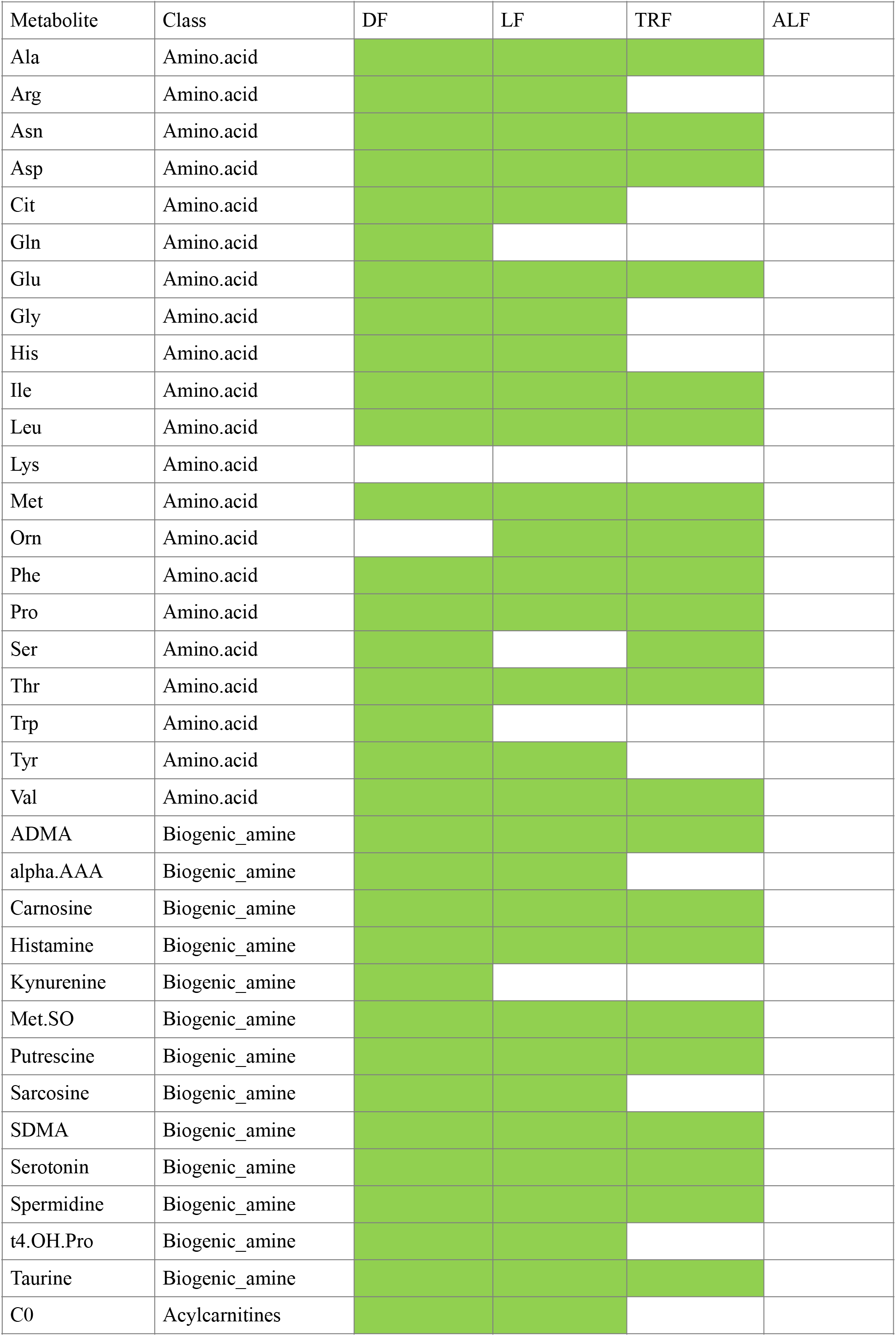
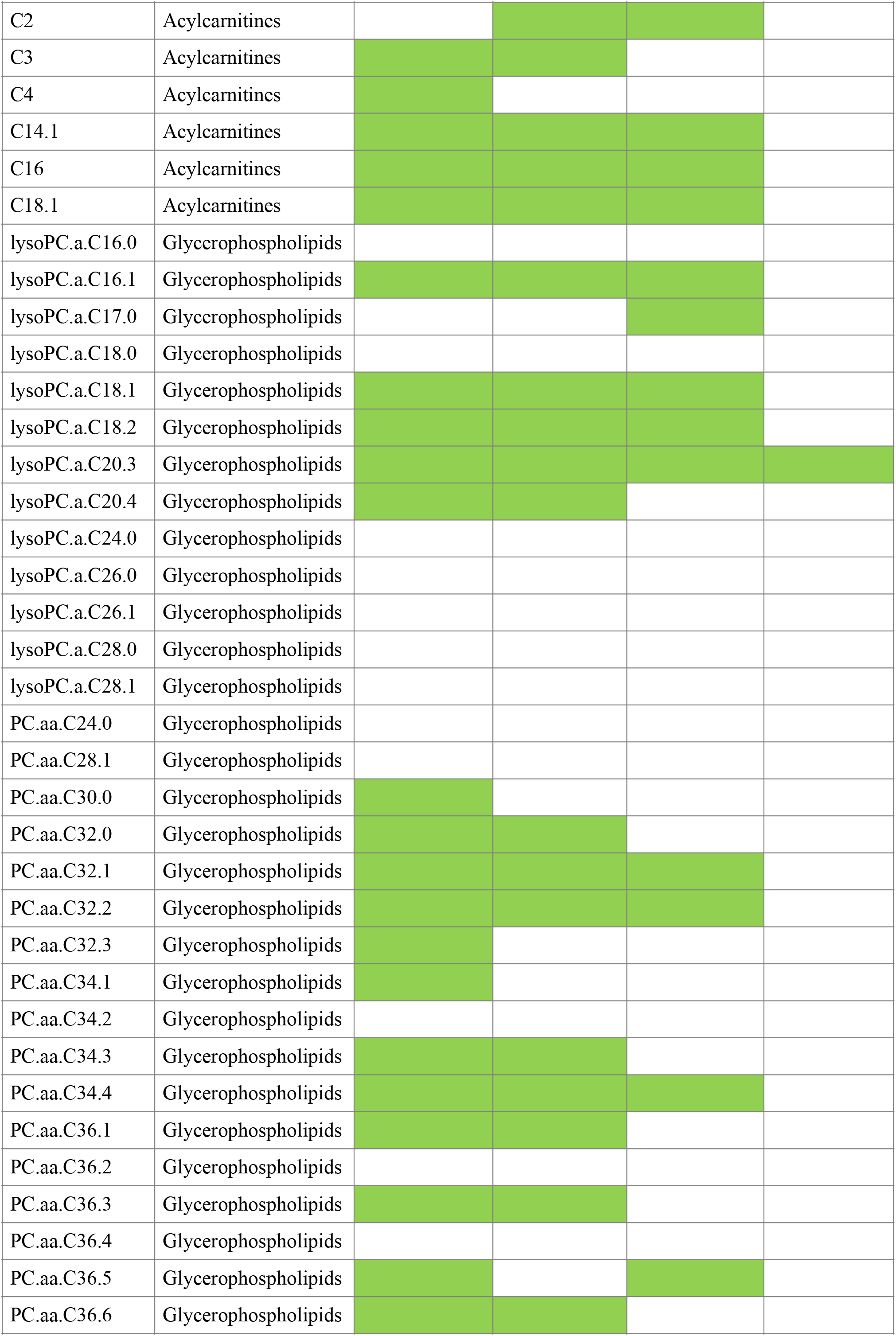
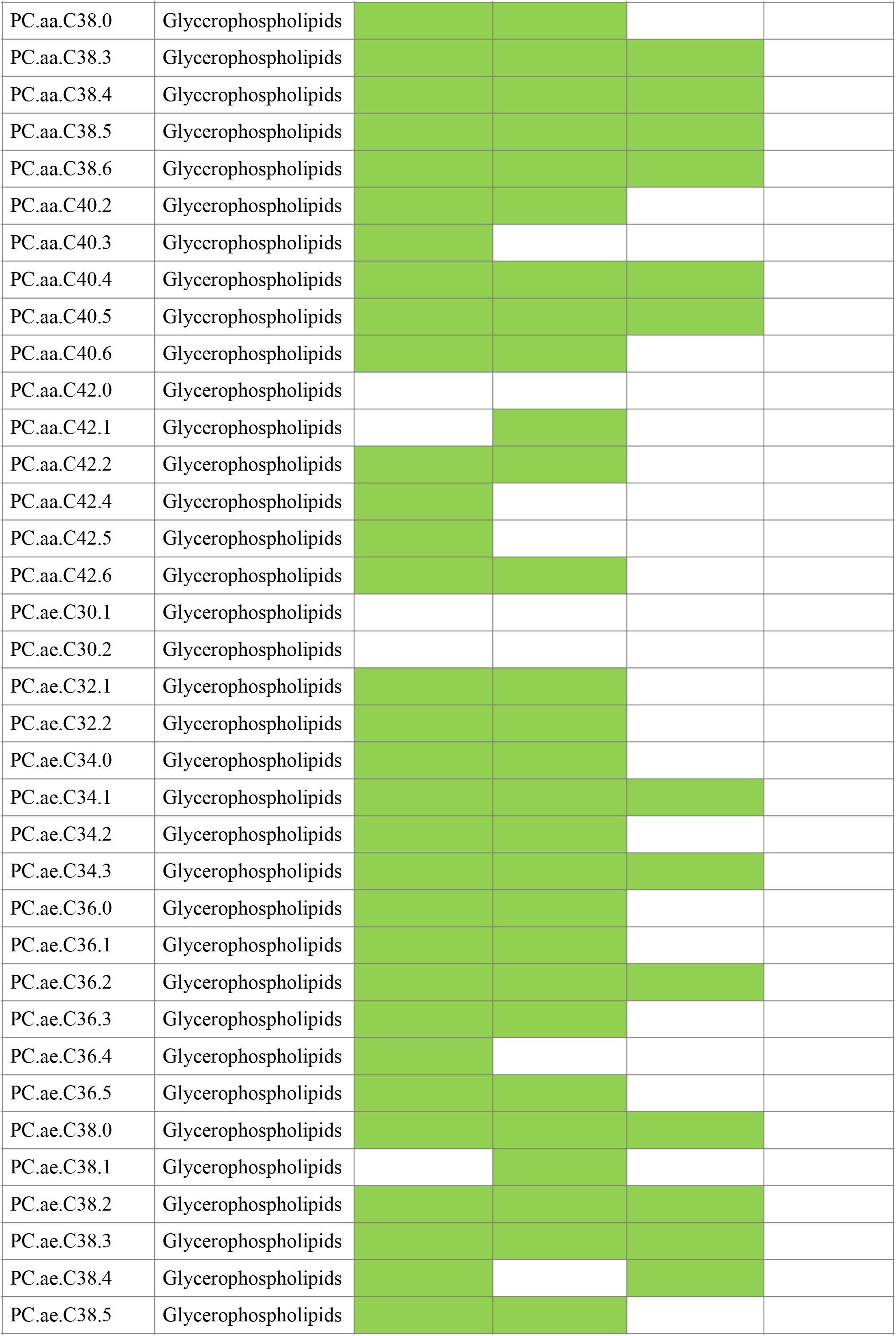
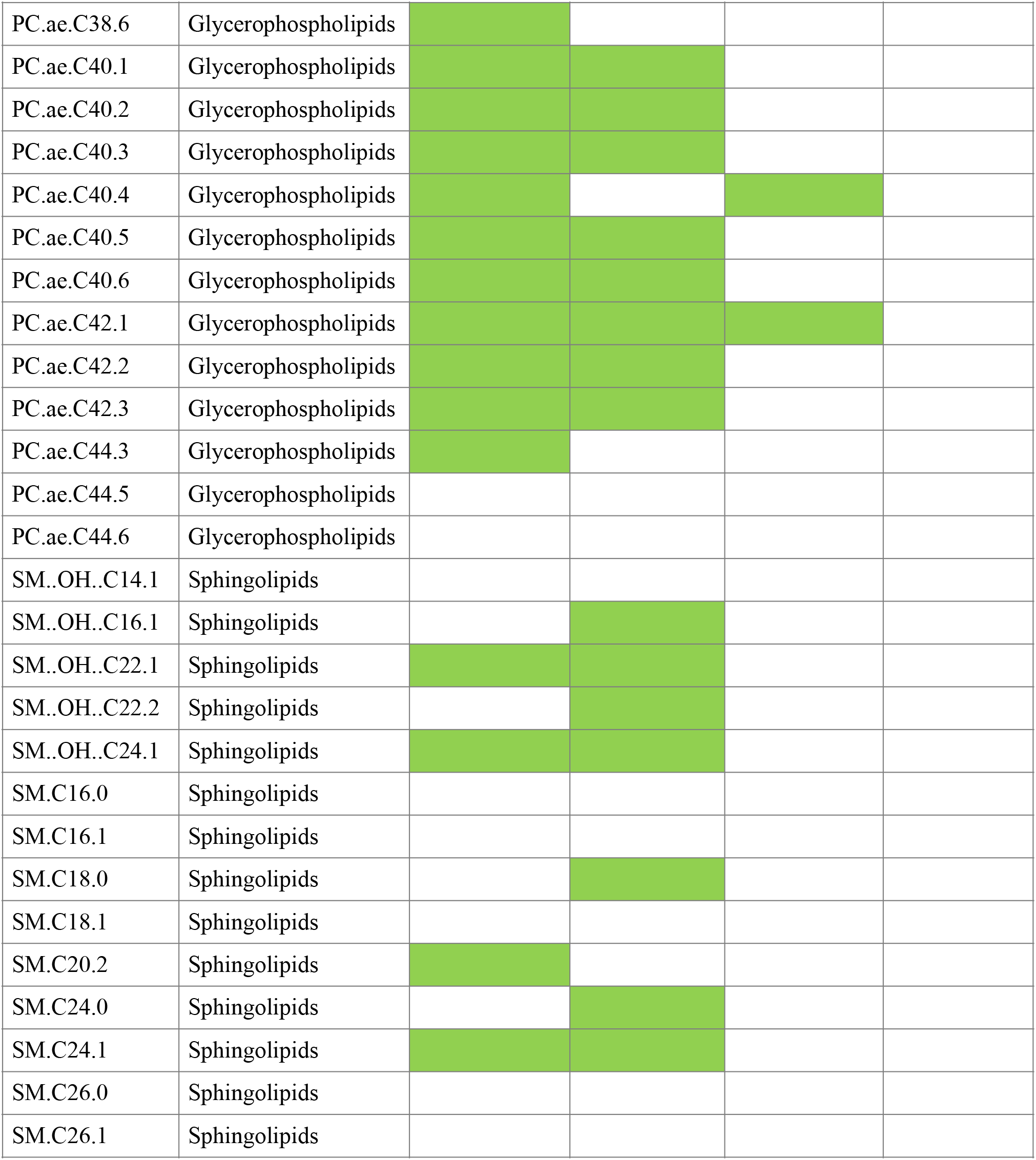
Highlighted in green are the metabolites rhythmic in each treatment group (according to the circadian programmes and ANOVA). DF: 101 y, 33 n; LF: 91 y, 43 n; TRF: 50 y, 84 n; ALF: 1 y, 133 n. Relates to Figs. 3 and 5.

**Supp Table 6.**
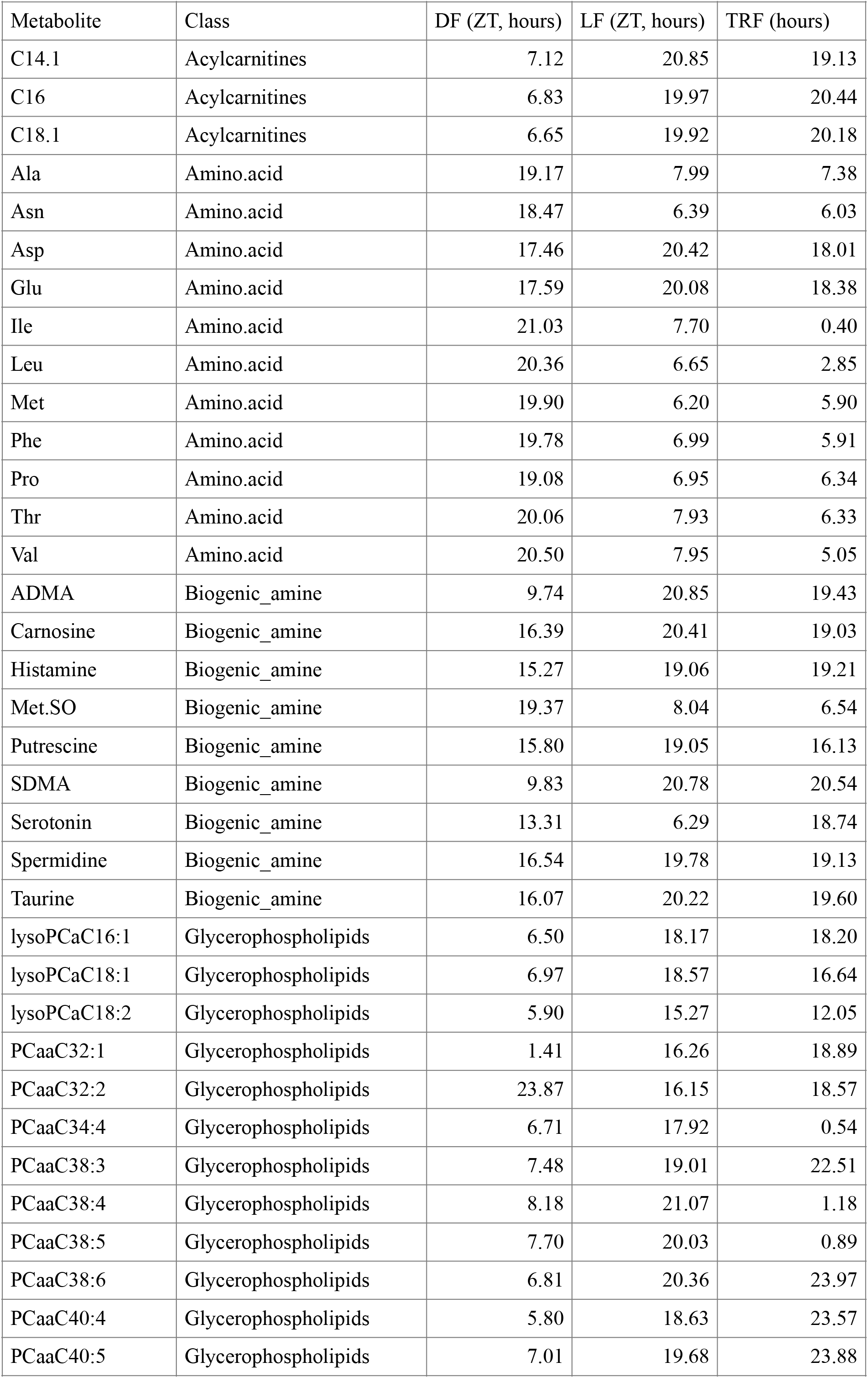
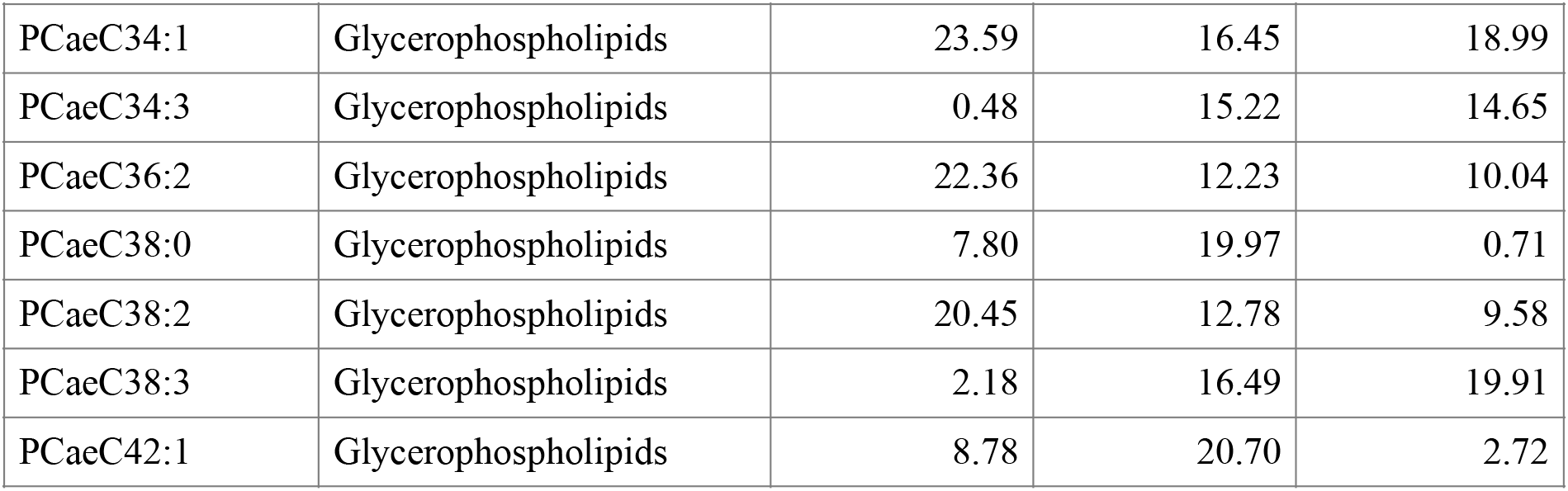
Timing of peak of each final candidate metabolite in the blood for each treatment group. For LF and TRF groups ZT 0/ 0 hours is the start of the feeding window and the time of lights on for LF, while for DF ZT0 is the start of the fasting window and the time of lights on. LF and DF groups are in 12h:12h light:dark although feeding in the day and night respectively. TRF are in constant darkness although feed for the same 12h window (0-12 hours) as the LF group (same experimenter time). Relates to Figs 3 and 5.

| Metabolite | Class | DF (ZT, hours) | LF (ZT, hours) | TRF (hours) |
| --- | --- | --- | --- | --- |
| C14.1 | Acylcarnitines | 7.12 | 20.85 | 19.13 |
| C16 | Acylcarnitines | 6.83 | 19.97 | 20.44 |
| C18.1 | Acylcarnitines | 6.65 | 19.92 | 20.18 |
| Ala | Amino.acid | 19.17 | 7.99 | 7.38 |
| Asn | Amino.acid | 18.47 | 6.39 | 6.03 |
| Asp | Amino.acid | 17.46 | 20.42 | 18.01 |
| Glu | Amino.acid | 17.59 | 20.08 | 18.38 |
| Ile | Amino.acid | 21.03 | 7.70 | 0.40 |
| Leu | Amino.acid | 20.36 | 6.65 | 2.85 |
| Met | Amino.acid | 19.90 | 6.20 | 5.90 |
| Phe | Amino.acid | 19.78 | 6.99 | 5.91 |
| Pro | Amino.acid | 19.08 | 6.95 | 6.34 |
| Thr | Amino.acid | 20.06 | 7.93 | 6.33 |
| Val | Amino.acid | 20.50 | 7.95 | 5.05 |
| ADMA | Biogenic_amine | 9.74 | 20.85 | 19.43 |
| Carnosine | Biogenic_amine | 16.39 | 20.41 | 19.03 |
| Histamine | Biogenic_amine | 15.27 | 19.06 | 19.21 |
| Met.SO | Biogenic_amine | 19.37 | 8.04 | 6.54 |
| Putrescine | Biogenic_amine | 15.80 | 19.05 | 16.13 |
| SDMA | Biogenic_amine | 9.83 | 20.78 | 20.54 |
| Serotonin | Biogenic_amine | 13.31 | 6.29 | 18.74 |
| Spermidine | Biogenic_amine | 16.54 | 19.78 | 19.13 |
| Taurine | Biogenic_amine | 16.07 | 20.22 | 19.60 |
| lysoPCaC16:1 | Glycerophospholipids | 6.50 | 18.17 | 18.20 |
| lysoPCaC18:1 | Glycerophospholipids | 6.97 | 18.57 | 16.64 |
| lysoPCaC18:2 | Glycerophospholipids | 5.90 | 15.27 | 12.05 |
| PCaaC32:1 | Glycerophospholipids | 1.41 | 16.26 | 18.89 |
| PCaaC32:2 | Glycerophospholipids | 23.87 | 16.15 | 18.57 |
| PCaaC34:4 | Glycerophospholipids | 6.71 | 17.92 | 0.54 |
| PCaaC38:3 | Glycerophospholipids | 7.48 | 19.01 | 22.51 |
| PCaaC38:4 | Glycerophospholipids | 8.18 | 21.07 | 1.18 |
| PCaaC38:5 | Glycerophospholipids | 7.70 | 20.03 | 0.89 |
| PCaaC38:6 | Glycerophospholipids | 6.81 | 20.36 | 23.97 |
| PCaaC40:4 | Glycerophospholipids | 5.80 | 18.63 | 23.57 |
| PCaaC40:5 | Glycerophospholipids | 7.01 | 19.68 | 23.88 |
| PCaeC34:1 | Glycerophospholipids | 23.59 | 16.45 | 18.99 |
| PCaeC34:3 | Glycerophospholipids | 0.48 | 15.22 | 14.65 |
| PCaeC36:2 | Glycerophospholipids | 22.36 | 12.23 | 10.04 |
| PCaeC38:0 | Glycerophospholipids | 7.80 | 19.97 | 0.71 |
| PCaeC38:2 | Glycerophospholipids | 20.45 | 12.78 | 9.58 |
| PCaeC38:3 | Glycerophospholipids | 2.18 | 16.49 | 19.91 |
| PCaeC42:1 | Glycerophospholipids | 8.78 | 20.70 | 2.72 |

**Supp Table 7.** Degrees of Freedom (df), log-Likelihood (logLik), AICc, ΔAICc (AICc_i_ – AICc_min_) and AICc w (AICc weight) for each linear model in the schizont/parasite proportion/density analysis ordered in descending fit (best-fitting model at the top). The response variable for each model is either schizont proportion (Schizont.prop), parasite density (Parasite.dens) or schizont density (Schizont.dens) and the random effect is “mouseID”. “Treatment” refers to the treatment group (DF, LF, TRF or ALF), “hours in culture” refers to the number of hours spent in culture since being extracted from the mice, and “hours since isoleucine” refers to the number of hours since isoleucine was added to the cultures. Treatment, hours in culture and hours since isoleucine were all fitted as factors. Relates to Fig 6.

| **Model description:** | **df** | **logLik** | **AICc** | **ΔAICc** | **AICc w** |
| --- | --- | --- | --- | --- | --- |
| **A) Schizont.prop ~ + (1\|mouseID)** |  |  |  |  |  |
| treatment | 4 | 106.26 | -203.8 | 0.00 | 0.996 |
| treatment + hours in culture | 7 | 104.29 | -192.6 | 11.26 | 0.004 |
| treatment * hours in culture | 10 | 98.81 | -173.5 | 30.37 | 0.000 |
| null | 3 | 80.96 | -155.5 | 48.31 | 0.000 |
| hours in culture | 6 | 74.89 | -136.3 | 67.54 | 0.000 |
| **B) Parasite.dens ~ + (1\|mouseID)** |  |  |  |  |  |
| null | 3 | 0.14 | 6.6 | 0.00 | 0.836 |
| treatment | 4 | -0.63 | 10.7 | 4.16 | 0.104 |
| hours in culture | 4 | -1.30 | 12.1 | 5.50 | 0.053 |
| treatment + hours in culture | 5 | -2.07 | 16.4 | 9.88 | 0.006 |
| treatment * hours in culture | 6 | -2.79 | 20.9 | 14.36 | 0.001 |
| **C) Schizont.dens ~ + (1\|mouseID)** |  |  |  |  |  |
| treatment + hours since isoleucine | 6 | 69.67 | -125.3 | 0.00 | 0.887 |
| hours since isoleucine | 4 | 65.04 | -121.2 | 4.14 | 0.112 |
| treatment * hours since isoleucine | 8 | 65.67 | -111.6 | 13.65 | 0.001 |
| null | 3 | 56.87 | -107.2 | 18.10 | 0.000 |
| treatment | 5 | 56.67 | -101.9 | 23.39 | 0.000 |
| **D) Parasite.dens ~ + (1\|mouseID)** |  |  |  |  |  |
| hours since isoleucine | 4 | 12.96 | -17.0 | 0.00 | 0.568 |
| null | 3 | 11.49 | -16.4 | 0.56 | 0.430 |
| treatment + hours since isoleucine | 6 | 9.10 | -4.1 | 12.85 | 0.001 |
| treatment | 5 | 7.76 | -4.1 | 12.90 | 0.001 |
| treatment * hours since isoleucine | 8 | 10.47 | -1.2 | 15.75 | 0.000 |

## Notes

### Competing Interest Statement

The authors have declared no competing interest.

